# Barcode Fusion Genetics-Protein-fragment Complementation Assay (BFG-PCA): tools and resources that expand the potential for binary protein interaction discovery

**DOI:** 10.1101/2021.07.27.453987

**Authors:** Daniel Evans-Yamamoto, François D. Rouleau, Piyush Nanda, Koji Makanae, Yin Liu, Philippe C. Després, Hitoshi Matsuo, Motoaki Seki, Alexandre K. Dube, Diana Ascencio, Nozomu Yachie, Christian R. Landry

**Author notes:** To whom correspondence should be addressed., Correspondence may also be addressed to Nozomu Yachie. [Piyush Nanda], Department of Molecular and Cellular Biology, Harvard University, Cambridge, Massachusetts, 02138, United States of America. [Yin Liu], School of Biomedical Engineering, University of British Columbia, Vancouver, British Columbia, V6T 1Z3, Canada. [Hitoshi Matsuo], Department of Biological Sciences, Graduate School of Science, The University of Tokyo, Tokyo, 113-0033, Japan. [Motoaki Seki], Department of Molecular Oncology, Graduate School of Medicine, Chiba University, Chiba, 260-8670, Japan. [Nozomu Yachie], School of Biomedical Engineering, University of British Columbia, Vancouver, British Columbia, V6T 1Z3, Canada.

## Abstract

Barcode fusion genetics (BFG) utilizes deep sequencing to improve the throughput of protein-protein interaction (PPI) screening in pools. BFG has been implemented in Yeast two-hybrid (Y2H) screens (BFG-Y2H). While Y2H requires test protein pairs to localize in the nucleus for reporter reconstruction, Dihydrofolate Reductase Protein-Fragment Complementation Assay (DHFR-PCA) allows proteins to localize in broader subcellular contexts and proves to be largely orthogonal to Y2H. Here, we implemented BFG to DHFR-PCA (BFG-PCA). This plasmid-based system can leverage ORF collections across model organisms to perform comparative analysis, unlike the original DHFR-PCA that requires yeast genomic integration. The scalability and quality of BFG-PCA were demonstrated by screening human and yeast interactions for >11,000 protein pairs. BFG-PCA showed high-sensitivity and high-specificity for capturing known interactions for both species. BFG-Y2H and BFG-PCA capture distinct sets of PPIs, which can partially be explained based on the domain orientation of the reporter tags. BFG-PCA is a high-throughput protein interaction technology to interrogate binary PPIs that exploits clone collections from any species of interest, expanding the scope of PPI assays.

## INTRODUCTION

In cellular systems, proteins form functional modules and/or complexes that underlie most biological processes by physically interacting with each other (1, 2). Discovering such interaction networks is one of the main goals of systems biology. Two major approaches to detect protein-protein interactions (PPIs) have contributed the bulk of the current data, affinity purification followed by mass spectrometry (AP/MS), and methods such as Yeast two-hybrid (Y2H) and protein complementation assay (PCA). The former approach detects biomolecular association among groups of proteins from cellular fractions (3–6), whereas the latter detects direct “binary” or pairwise PPIs, by tagging each interaction partner, the bait and the prey, using reporter protein fragments (7–9). Other approaches such as proximity-dependent biotinylation *in vivo* (5, 10, 11), co-elution and co-fractionation (12, 13) and protein-cross linking (14, 15) also contribute to the dissection of PPI networks, with varying degrees of resolution.

Binary interaction screenings are powerful approaches owing to their relatively simple implementation in terms of instrumentation. Up to now, systematic high-quality Y2H screening (16) has revealed the largest binary interactome network to date, covering the entire human and yeast proteomes (17–19). Because of its scalability, Y2H has also been applied to a large number of model organisms, including for instance Arabidopsis and Drosophila (20–23). Despite such efforts, we are far from a complete interactome map when considering various “proteoforms” (24, 25), disease mutations (26) and protein polymorphisms that can have distinct biophysical interaction profiles (27).

One of the limiting factors associated with binary detection methods is the need to perform pairwise tests between baits and preys in a comprehensive manner, and these pairwise tests are dominated by negative results, i.e. most protein pairs do not interact. However, the application of next generation sequencing (NGS) has played a key role in increasing throughput, and thus, interactome coverage (18, 28– 36). Combined with methods that involve cell survival as detection signals, NGS facilitates the exploration of the search space of PPIs because of the enrichment of positive PPIs. Early efforts have screened for interactors of a given bait in a pool of prey proteins (one against all), which were identified by sequencing of the prey ORF region after selection (30, 31). Further studies that have implemented a pooled screening approach using NGS, Stitch-seq, allowed the identification of both bait and prey as pair information in pooled assays through fusion PCR of bait and prey ORFs after selecting for interacting pairs (18, 28). Several other approaches exploiting this principle have been implemented. They include Barcode Fusion Genetics (BFG) (32), PPi-seq (34, 37), which use synthetic DNA barcodes to tag gene of interests and CrY2H-seq (33), rec-YnH (36), and RLL-Y2H (35, 36), which use the Open Reading Frame (ORF) sequences themselves to identify protein pairs. Using ORF sequences as identifiers offers simplicity to the design but DNA barcodes may be more reliable in terms of accuracy and performance, and may reduce sequencing costs, although they may require more investment upstream of the screens.

BFG was recently adapted to Y2H screening. In BFG experiments, bait and prey plasmids contain DNA barcodes that are fused through intra-cellular recombination in cells that survive selection on a specific media. Sequencing of the fused barcodes allows the identification of the interacting pairs in bulk (**Figure 1A**). Because the barcode fusion technology is portable to other approaches in yeast genetics, it could be used to adapt other binary mapping methods to pooled screening and thus enable a better coverage of PPI networks. Indeed, different assays have little but significant overlap of positive interactions, and it is important to assay PPIs with multiple orthogonal assays to comprehensively map interactomes (38, 39). For instance, systematic benchmarking of various complementary assays in yeast and human cells has reported that each method captures only ~35% of the confident positive reference PPI set (HsPRSv1) (38). Even more revealing for this study, for binary assays, the currently reported *S. cerevisiae* PPIs by Y2H and PCA share only 525 unique interactions (Y2H: 12,995; PCA: 6,739; Union: 19,209; Jaccard Index: 0.027), despite each method having proteome wide PPI mapping efforts of similar quality (9, 19, 40). There are many reasons why different methods cover different parts of PPI networks. For instance, reporter proteins or protein fragments are fused at either the N or C termini and some may require the localization of proteins to specific cell compartments (41) **(Figure 1B and C)**.

**Figure 1.**
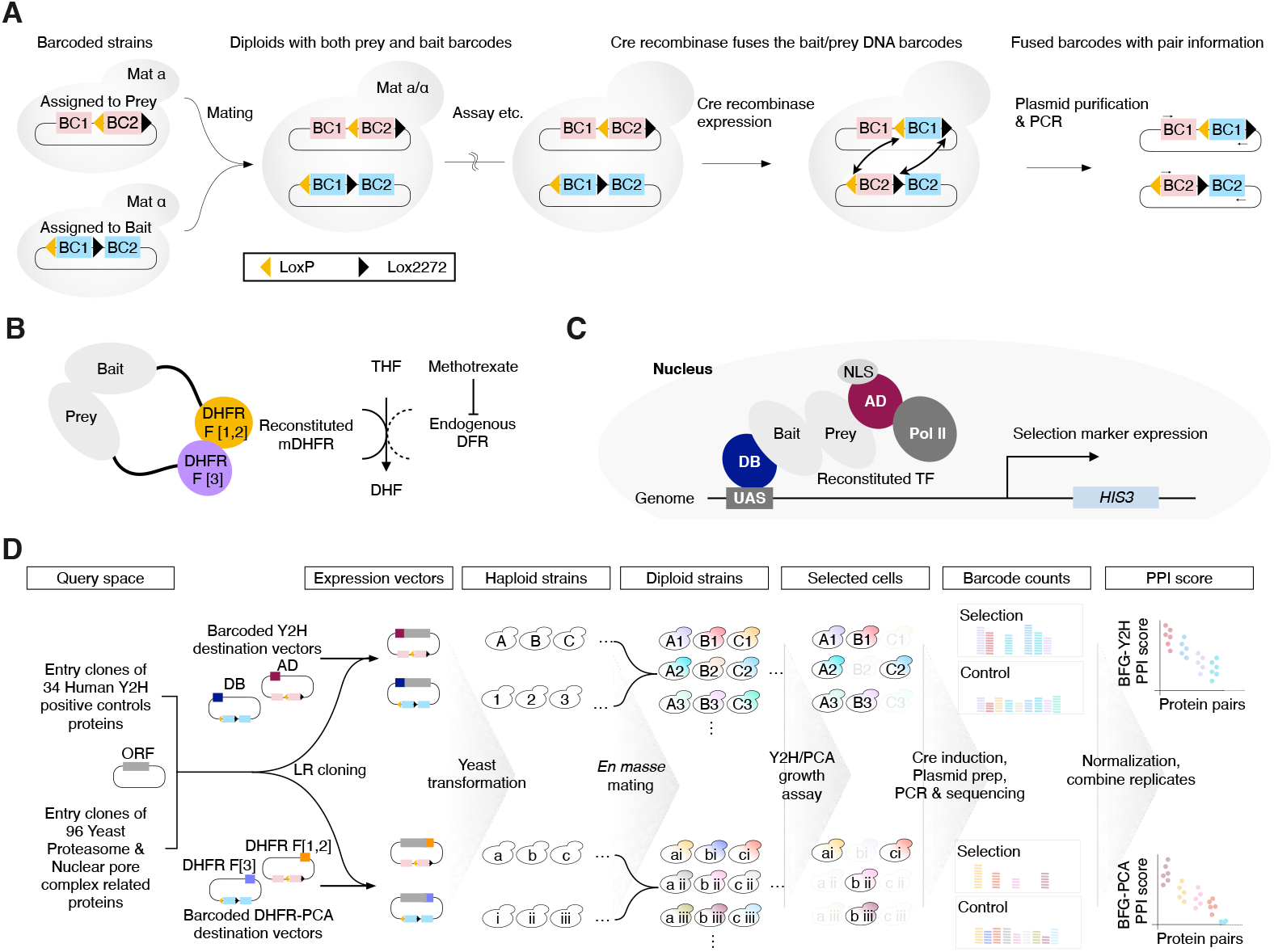
Overview of this study. (**A**) Schematic of Barcode Fusion Genetics (27). BFG barcode cassettes are assigned to each of the bait and prey with 2 DNA barcodes (BC1 and BC2), with one of them flanked by Cre recombination site LoxP and Lox2272. The bait and prey plasmids with BFG barcode cassettes are introduced to MATα and MATa cells, respectively. Upon mating, the diploid cells have both the bait and prey plasmids and their barcode cassettes. By inducing expression of the Cre recombinase, the BC1bait and BC2_prey_ are swapped between the plasmids, resulting in barcode fusions BC1_prey_-BC1_bait_ and BC2_bait_-BC2_prey_, having the bait-prey pair information. Each of the bait and prey barcodes have a common primer site flanking the barcode region unique to the type of barcode, enabling specific PCR amplification of the BC1_prey_-BC1_bait_ and BC2_prey_-BC2_bait_ products. By counting the number of barcode pairs by deep sequencing, one can estimate the relative abundance of diploids in the pool. (**B**) Illustration of the DHFR-PCA reporter. In DHFR-PCA, DHFR F[1,2] and DHFR F[3] are fused to bait and prey proteins, respectively. Upon interaction, the DHFR fragments come in proximity, reconstituting the methotrexate-resistant murine DHFR enzyme (mDHFR) while the conditionally essential endogenous DHFR is inhibited by the drug methotrexate (MTX). (**C**) Illustration of the Y2H reporter. In Y2H, the DNA binding domain (DB) and the activator domain (AD) of the Gal4 transcription factor (TF) are fused to bait and prey proteins, respectively. The fused proteins are localized to the nucleus by the nuclear localization signal (NLS). The DB domain will bind to the upstream activation sequence (UAS). When the bait and prey proteins interact, the Gal4TF is reconstituted, recruiting RNA polymerase II, expressing the selection marker of choice. We used the *HIS3* marker with medium lacking histidine for Y2H selection throughout this study. (**D**) Overview of the BFG screening. We queried 115 proteins from Human and Yeast, and Gateway cloned them to barcoded Y2H and DHFR-PCA destination vectors with 2 barcode replicates each. We individually transformed haploid strains with the barcoded expression vectors. The haploid strains were pooled for *en masse* mating, generating all possible bait-prey pairs of diploids. After selecting diploid cells, we performed pooled selection for each method. After selection, we induced Cre expression for barcode fusion, purified the plasmids, and PCR amplified the barcodes for illumina sequencing. We counted barcodes, normalized them by the barcode counts in the control condition and background auto-activity of the strains. The replicates were combined for each protein pair, generating the final PPI score for each method to call PPIs.

BFG enables pooled matrix screening (all baits against all preys) that exploits various selection markers affecting growth (32). DHFR-PCA is a binary PPI detection method based on growth via the reconstitution of an engineered DHFR in yeast cells, which provides resistance to the drug methotrexate (9). Contrary to Y2H, DHFR-PCA does not require the addition of a nuclear localization for reporter activation, and in principle enables PPI detection in the protein pair’s native subcellular context. Until now, efforts to map PPIs by DHFR-PCA have focused on interactions present *in vivo* by tagging DHFR fragments at genomic loci, even when barcodes are used for pooled based assays (29, 34). Although this comes with many advantages, it also comes with limitations depending on the questions being addressed. For instance, protein expression levels are largely regulated by the environment, making interactions of weakly expressed/unexpressed proteins difficult to detect in some conditions, including the ones used for testing. Having bait and prey proteins expressed from plasmids could help alleviate this limitation. Controlling or uniformalizing expression level may help differentiate transcriptional versus post transcriptional effects on PPIs in experiments comparing different growth conditions (34, 42). Another advantage of plasmid-based screening is that it allows for screening PPIs for protein variants, or among proteins from other species or between species, provided the coding sequences can be cloned and expressed in yeast. Here, we developed and made publicly available affordable resources for BFG-DHFR-PCA (henceforth BFG-PCA) and we used this resource to demonstrate the efficacy of BFG-PCA by screening 11,232 bait-prey pairs (**Figure 1D**). We show that BFG-PCA enables the detection of *in vivo* PPIs and the comparison side-byside with BFG-Y2H demonstrates that they capture distinct sets of PPIs.

## MATERIAL AND METHODS

### DNA oligomers

All PCR primers and gene fragments used in this study are listed in **Supplementary Table S1**.

### Reagents

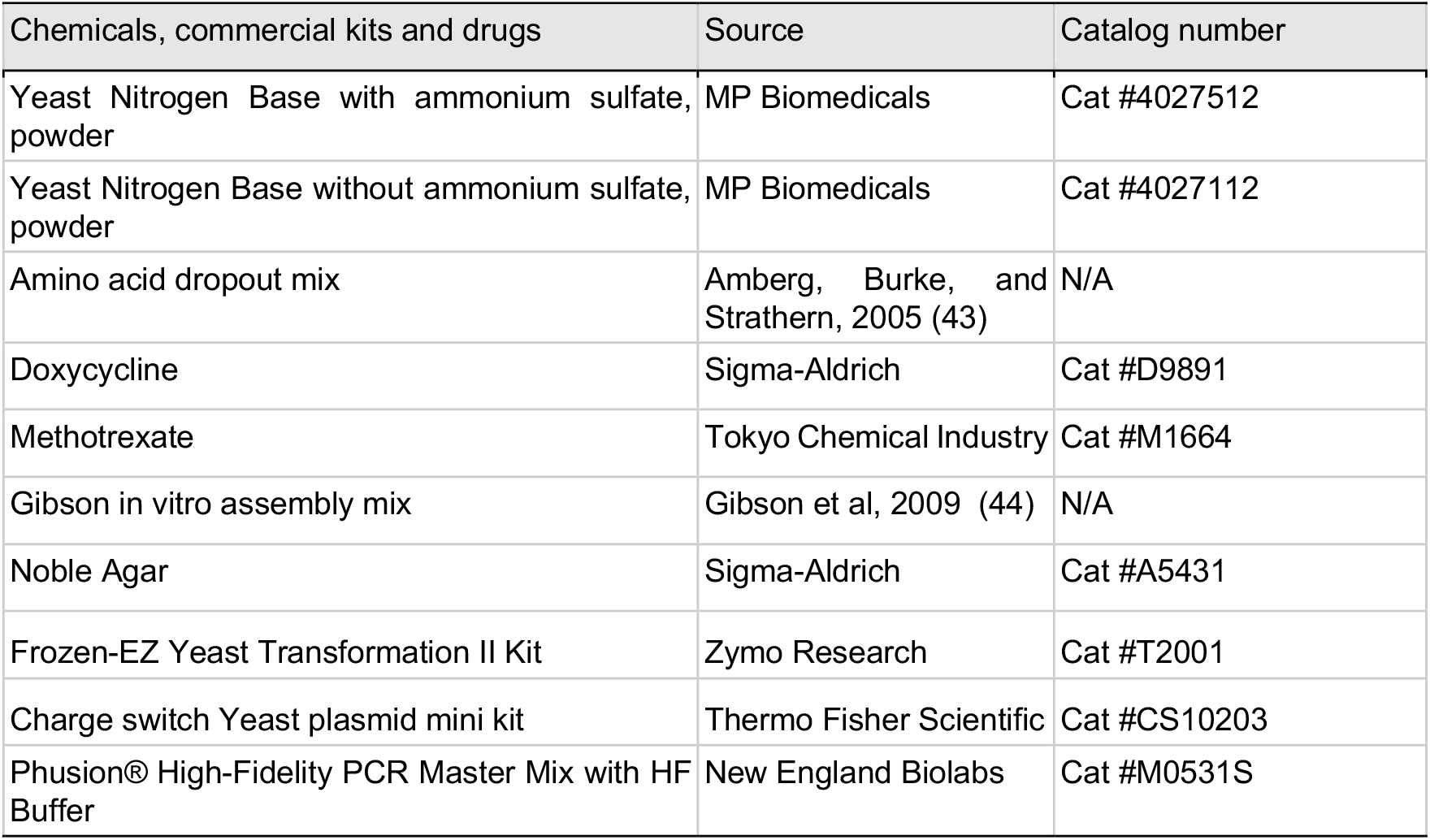

### Biological resources

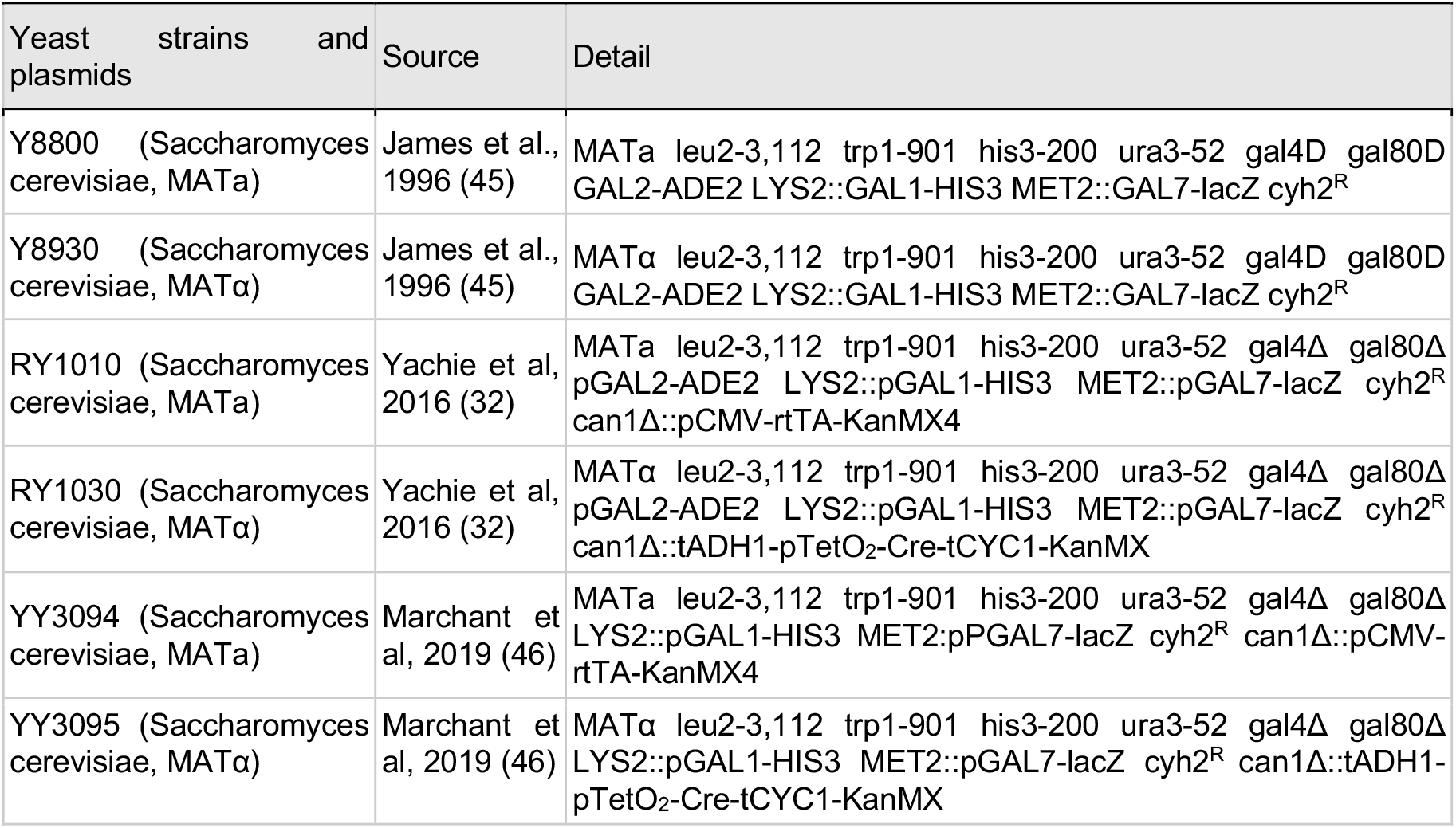
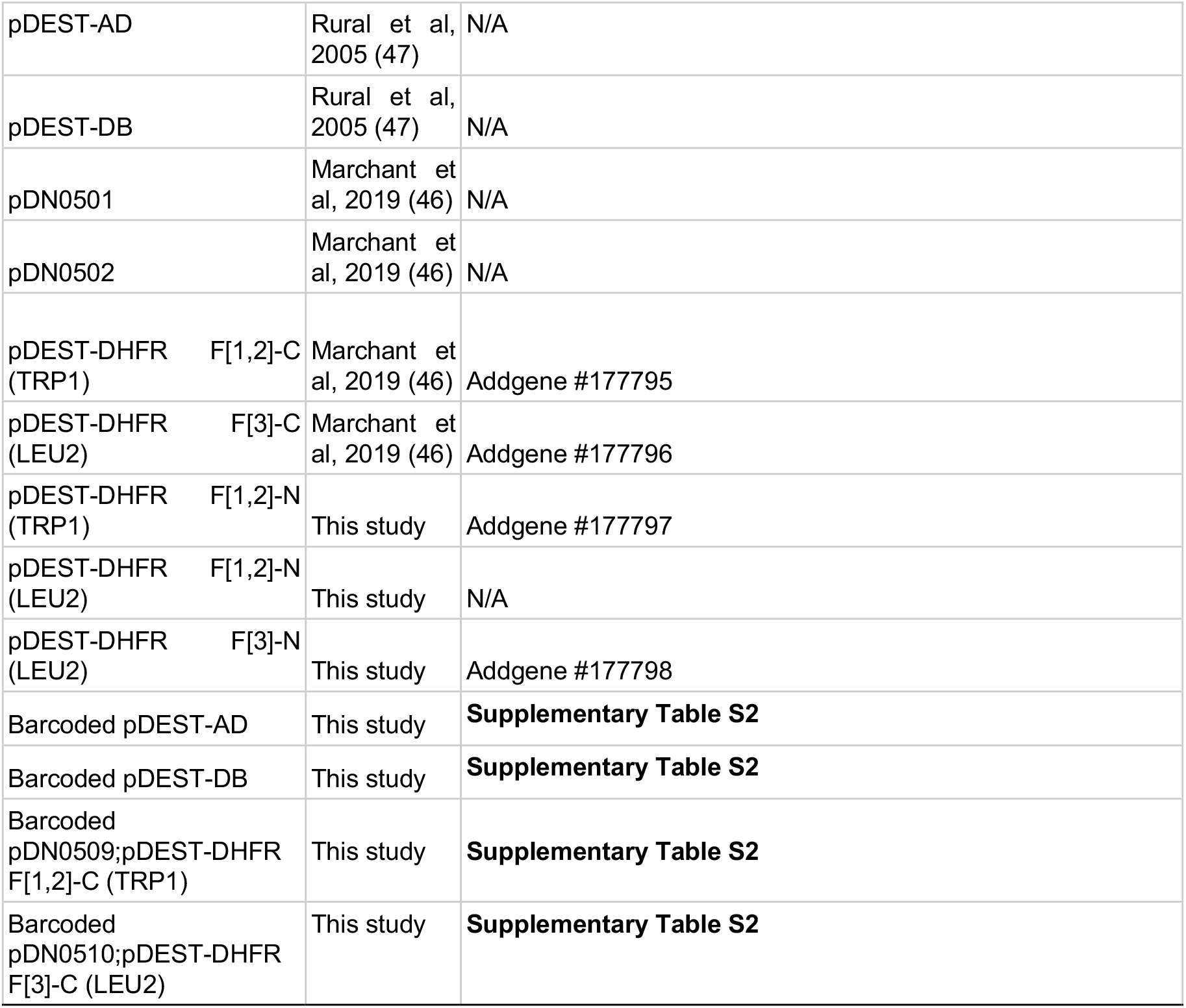

### Computational resources

| Software | Source |
| --- | --- |
| Python version 3.6.1 | https://www.python.org |
| R version 4.0.4 | https://www.r-project.org |
| BLASTn version 2.4.0 | Altschul et al, 1990 (48) |
| PDBePISA version 1.52 | Krissinel and Henrick, 2007 (49) |
| Pymol version 2.5.0 | Schrödinger Inc. |

### Plasmid construction

For the plasmid-based DHFR-PCA and BFG-PCA, we used Gateway cloning-compatible plasmid vectors which we previously constructed based on the Y2H plasmids pDEST-AD and pDEST-DB (46). Plasmid-based DHFR-PCA vectors bearing the DHFR fragment domain on the N-terminus end of the protein (pDEST-DHFR F[1,2]-N and pDEST-DHFR F[3]-N) were constructed for this study. To generate pDEST-DHFR F[1,2]-N (LEU2) and pDEST-DHFR F[3]-N (LEU2), the DB domain of pDN0502 (LEU2) was replaced with DHFR fragments by ligation. The backbone fragment was prepared by restriction digestion of pDN0502 using HindIII and NotI and purified by size-selection on gel. The insert DHFR fragments DEY030 (DHFR F[1,2]) and DEY031 (DHFR F[3]) were ordered as gene fragments (TWIST biosciences). The fragments were amplified using primers DEY032 and DEY033. The insert fragments were purified on gel after digestion with HindIII and NotI. The pDN0502 backbone and each of the inserts were used for ligation to generate pDEST-DHFR F[1,2]-N (LEU2) and pDEST-DHFR F[3]-N (LEU2). To generate pDEST-DHFR F[1,2]-N (TRP1), pDN0501 and pDEST-DHFR F[1,2]-N (LEU2) were digested with I-CeuI and I-SceI to size select the backbone and insert, respectively. The two fragments were used for ligation to generate the pDEST-DHFR F[1,2]-N (TRP1) plasmid. After Gateway LR cloning of entry clones to these destination plasmids, the expression plasmids encode DHFR-linker-protein fusion protein with the following linker sequence GGGSTSTSLYKKVG. The plasmids were each confirmed for their correct construction by Sanger sequencing.

### Construction of Y2H and DHFR plasmids and strains for PPI assay

Expression plasmids were generated by subcloning ORF regions of entry plasmids to destination plasmids by Gateway LR reaction (50). In detail, 10 ng each of entry plasmid and destination plasmid was mixed with Gateway LR clonase II (Invitrogen) in a 2 μL final volume and incubated at 25 °C for at least 16 hours. The entire volume of the enzymatic reaction was used to transform 25 μL of NEB5-alpha chemically competent *E.coli* cells, prepared as previously described (46). The transformation was performed as in (51) but with selection on LB+ampicillin plates followed by incubation overnight at 37°C instead of direct inoculation to liquid culture. The colonies were scraped and cultured in 5 mL LB+ampicillin for plasmid purification.

### Yeast medium for Y2H and DHFR-PCA assays

Haploid and diploid strain cultures of Y2H and PCA samples were cultured in SC-Leu+Ade+His/SC-Trp+Ade+His and SC-Leu-Trp+Ade+His, respectively. Mating was performed in YPAD medium. For Y2H selection, the control condition was on SC-Leu-Trp+0.18 mg /mL Ade+8 mM His plates, and selection conditions was on SC-Leu-Trp-His+0.18 mg/mL Ade and SC-Leu-Trp-His+ 0.18 mg /mL Ade+ 1 mM amino-l,2,4-triazole (3-AT). Preparation was carried out as previously described (51), and shown in **Supplementary Note S2**. The amino acid dropout mix (DO mix) was prepared as previously described (43).

For DHFR-PCA and BFG-PCA screenings, methotrexate (MTX) (Tokyo Chemical Industry co., ltd.) was dissolved in 20 mL of DMSO according to the final concentration. The control condition was SC-Leu-Trp-Ade+8 mM His+2.0 % (v/v) DMSO, and default selection was SC-Leu-Trp-Ade+8 mM His+2.0 % (v/v) DMSO+ 200 μg/mL methotrexate. Preparation was carried out as previously described (46), and shown in **Supplementary Note S2**. The 10× DO mix solution was prepared by dissolving 15 g of the powder DO mix in deionized water, and filtered for sterilization.

### DHFR-PCA spot and pintool assays

The purified pDHFR F[1,2]-ORF and pDHFR F[3] expression plasmids were used to transform stains YY3094 and YY3095, respectively. The pDHFR F[3]-ORF (bait) and pDHFR F[1,2]-ORF (prey) transformants were respectively selected on SC-Leu+Ade+His or SC-T rp+Ade+His plates at 30 °C for 48 hours. The resulting haploid bait and prey strains were pre-cultured individually in 96-well plates with 200 μL of media and incubated at 30 °C for 2 overnights. SC-Leu+Ade+His and SC-Trp+Ade+His media were used to culture bait and prey strains, respectively. The haploid strains were mated by spotting 5 μl each of the bait and prey culture for all protein pair combinations on YPAD plates and incubated overnight at 30 °C for mating. The mated samples were inoculated to 1 mL of SC-Leu-Trp+Ade+His liquid medium in a deep 96-well plate, and diploid cells were selected by incubation at 30 °C with 200 rpm agitation overnight. The resulting diploid culture was centrifuged at 500×g and resuspended in autoclaved Millipore quality H_2_O twice and subjected to OD_600_nm measurement. For selection, the samples were spotted at OD_600_nm = 0.5 at a volume of 5 μl on 3.0% (w/v) agar plates of SC-Ade-Leu-Trp+2% (v/v) DMSO (-MTX), and SC-Ade-Leu-Trp+2% (v/v) DMSO + 200 μg/mL methotrexate ( + MTX). The selection plates were incubated for 72 hours at 30 °C for growth scoring, and further incubated at 30 °C and observed every 24 hours.

The pintool assay was performed using the generated DHFR-PCA strains based on standard procedures previously described (52).

### Barcode Fusion assay

Barcoded plasmids were transformed each into Y2H (Y8800 and Y8930), BFG-Y2H (RY1010 and RY1030), and BFG-PCA (YY3094 and YY3095) strains, mated, and selected for diploid as above. The diploid samples were subjected to doxycycline induction after adjusting OD_600_nm to 1.0 in 2.5mL of SC-Leu-Trp+Ade+10 μg/mL doxycycline, and incubated 30 °C in rotation for one overnight until the OD_600_nm reached 5.0. The samples were lysed as previously described (53), and genotyping PCR was performed with conditions as in Yachie *et al* (32) to check the BFG performance (**Supplementary Note S1 and Figure S1**).

### Spot assay to observe effects of DNA barcodes

For each of the pDEST-DHFR F[1,2]-C (TRP1) and pDEST-DHFR F[3]-C (LEU2), we randomly chose 2 unique barcoded destination plasmids (see below for barcoding procedures). For each ORF used in the assay, we constructed 3 expression plasmids by Gateway LR cloning the entry clone of the ORF to destination plasmids with and without barcodes. These barcoded expression plasmids were transformed each into BFG-PCA (YY3094 and YY3095) strains, mated, and selected for diploid as above. The diploid samples were inoculated in 3mL SC-Leu-Trp+Ade+His liquid medium and incubated at 30°C for two overnights. The resulting diploid culture was washed twice and subjected for selection as described above.

### Generation of barcoded Y2H and DHFR-PCA destination plasmid libraries

In total, 1,867 barcoded Y2H destination plasmids (1,137 for pDEST-AD and 730 for pDEST-DB) were generated as previously described (32). Briefly, two PCR products each having a random 25-bp flanked by lox sites and overlapping sequences were integrated into the SacI site of pDEST-AD or pDEST-DB via *in vitro* DNA assembly (44). This barcoded destination vector pool was transformed into One Shot ccdB Survival 2 T1^R^ Competent Cells (Invitrogen) that were spread on 245 mm × 245 mm square LB+ampicillin plates and incubated overnight at 37°C for colony isolation. Single colonies were picked by the QPix 450 robot (Molecular Device) and arrayed into a 384-well format. Two Row Column Plate-PCRs (RCP-PCRs) (32) were performed to identify clonal samples with their barcode sequences (BC-RCP-PCR) and to check the integrity of loxP and lox2272 sequences (Lox-RCP-PCR). RCP-PCR samples were multiplexed with other libraries, and sequenced on an Illumina MiSeq (2×250 bp paired-end sequencing). Pair of barcodes that had less than 5 % abundance within the well were eliminated to cancel out sequencing errors. The quality criteria was set so that only wells containing a single pair of barcode sequences with the designed elements were used for downstream processes.

The barcoded DHFR-PCA destination plasmids were generated similarly using destination plasmids pDEST-DHFR F[1,2]-C (TRP1) and pDEST-DHFR F[3]-C (LEU2). In total, 1,483 barcoded DHFR-PCA destination plasmids (893 for pDEST-DHFR F[1,2]-C and 590 for pDEST-DHFR F[3]-C) were generated. The destination vectors were digested with PI-PspI, and the two PCR products with random barcodes were inserted via *in vitro* DNA assembly (44). The PCR primers to generate the barcodes were altered from that of BFG-Y2H due to change in insert site. The primers used here are shown in **Supplementary Table S1**. The isolated bacterial colonies having barcoded destination vectors were prepared in the same manner as Y2H destination vectors. Two RCP-PCR were performed with the same design as in Y2H, but with minor modification in the primer used for Lox-RCP-PCR (**Supplementary Table S1**). The list of prepared barcodes are shown in **Supplementary Table S2**.

### Selection of ORFs used in this study

Positive controls were picked based on known Y2H interactions reported in the BioGRID database and retrieved from the CCSB human ORFeome resource (47). Nuclear pore complex (NPC) and proteasome related proteins were searched in the Uniprot (54) using keywords, “nuclear pore complex” and “proteasome”, respectively. Among the list, we accessed clones available from the *S.cerevisiae* Movable ORF collection (55), and quality controlled by Sanger sequencing using primer DEY034. The complete list of selected ORFs are shown in **Supplementary Table S3**.

### BFG-Y2H and BFG-PCA ready yeast strain generation

Barcoded expression plasmids with defined ORF-barcode associations were generated by one-by-one Gateway cloning. Similar to the non-barcoded expression vector preparation, ORF regions of entry plasmids were subclonded by Gateway LR cloning to a mix of 2 pre-assigned uniquely barcoded destination vectors. In detail, 10 ng of each entry plasmid and destination plasmid was mixed with Gateway LR clonase II (Invitrogen) in a 4 μL volume and incubated at 25 °C for at least 16 hours. Transformation of LR samples was performed in the same manner as non-barcoded samples. More than 5 colonies were scraped per sample to ensure representation of both barcodes, and cultured in 5 mL LB+ampicillin for plasmid purification. Purified plasmid was used to transform corresponding strains with appropriate selection medium. All prepared strains are listed in **Supplementary Table S3**.

### BFG-Y2H and BFG-PCA screenings

Haploid bait and prey strains were cultured to saturation by incubating at 30 °C in a static manner for approximately 60 hours in a 96-well deep well plate sealed with a breathable seal (Corning, BF-400-S). Each well contained 1 mL of SC-Leu+Ade+His or SC-Trp+Ade+His liquid media depending on the plasmids. Strains were pooled (AD,DB, DHFR F[1,2], or DHFR F[3]) by mixing 1 mL of 2 OD_600_nm equivalent cells for each strain. For mating, two groups of cell pools were mixed at equal amounts, and incubated at room temperature for 3 hours. After the incubation, the sample was spun down at 500×g for 4 minutes and then the cell pellet was spread on a YPAD plate. The plate was incubated at room temperature for 16 hours. The mated sample was scraped with autoclaved Millipore quality H_2_O, and then washed twice by spinning down the sample at 500×g for 4 minutes, and resuspending in SC-Leu-Trp+Ade liquid medium. The diploid cells were selected at a starting OD_600_nm of 1.0 in a 2 L flask containing 500 mL of SC-Leu-Trp+Ade liquid medium, and incubated for two overnights at 30 °C, 160 rpm. Fifty mL of the diploid selected culture sample was spun down, and washed twice with water. The screening was performed by plating an equivalent of 1 mL of 5.0 OD_600_nm cells per plate on 15 plates for each selection condition tested. The number of samples plated for selection was determined by a Monte-Carlo simulation model described in Yachie *et al* to ensure 100% of the positive diploid strain having at least 100 cells passing to the following step. The selected samples were collected after 72 hours of incubation at 30 °C. The collected samples was subjected to doxycycline induction after adjusting OD_600_nm to 1.0 in 25mL of SC-Leu-Trp+Ade+10 μg/mL doxycycline, and incubated 30 °C for one overnight until the OD_600_nm reached 5.0. The DNA extraction and deep sequencing library preparation was performed according to procedures shown in Yachie *et al*. Deep sequencing libraries were multiplexed with other libraries, and sequenced by Illumina MiSeq (2×250 bp paired-end sequencing). Reads were demultiplexed and fused DNA barcodes were counted by alignment of primer sequences and DNA barcodes using BLASTn version 2.4.0 (48) with the blastn-short option and an E-value threshold of 1e-10.

### BFG-Y2H and BFG-PCA data normalization

Both data normalization for BFG-Y2H and BFG-PCA data was performed with custom Python scripts. For BFG-Y2H data normalization, the procedures followed the method previously described (32). The detailed procedure for normalizing BFG-PCA data is described below.

For each condition and barcode fusion type (BC1-BC1 or BC2-BC2 fusion), the relative abundance of each diploid strain was estimated from the aggregated barcode count data. Note that a constant value of 1 was added to the barcode count of each strain to reduce noise for smaller values. For the non-selective (control) condition, all diploid strains are expected to grow, which results in high sequence complexity. Given that the deep sequencing depth is limited for the entire dynamic range of this complex pool, we first estimate the relative abundance computed as frequency for each *Bait_i_* or *Prey_j_* amongst all diploid strains *Dip_ij_* as

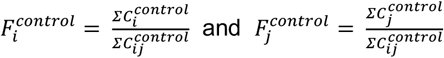

where *C* is the sequencing read count within each condition or barcode fusion type, respectively. Since chances for mating of each haploid combination is dependent on the relative abundance of each haploid strain, the frequency of diploid *Dip_ij_* (having *Bait_i_*, and *Prey_j_*) in non-selection condition (*F_ij_^control^*) can be estimated as

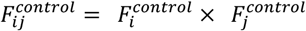

Relative growth of diploid in selection condition *F_ij_^selection^* was directly computed from raw count data as

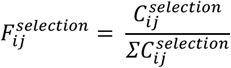

due to the sparse nature of PPI positives.

Based on relative abundance on non-selective and selective conditions, enrichment signal *s* was computed as

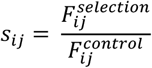

where *s* represents a degree of growth enrichment in favor of the selective condition for each diploid strain. Similar to the BFG-Y2H data, we observed different background levels of *s* for each haploid strain in BFG-PCA. We defined the background as the median of all *s* values for each haploid. The normalized score *ds* was computed for each diploid as

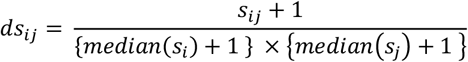

and subjected to PPI calling and analysis.

### PPI analysis

PPI analysis was performed by first aggregating all PPI scores for each protein pair, combining replicates and both bait-prey orientations tested. For each protein pair, PPI scores were sorted, and various percentiles (1, 5, 10, 15, 20,… 90, 95, 99), average, and median values were calculated. We ranked protein pairs based on each of the scoring methods (average, median, and each of the percentiles). Based on each scoring method, the protein pairs were sorted from highest to lowest, and subjected to computing Matthews Correlation Coefficient (MCC) (56) against the BioGRID database (version 4.4.198) (40) for quality assessment. We defined all PPIs reported in BioGRID by binary PPI detection methods (Y2H, PCA, Biochemical activity, Affinity Capture-Luminescence, Reconstituted Complex, Co-crystal Structure, and FRET) as positives, and categorized true positives (TP), true negatives (TN), false positives (FP), and false negatives (FN) for each rank threshold. We note that reports in BioGRID by PCA are not limited to DHFR-PCA. For each scoring method, the MCC for each rank threshold was computed as

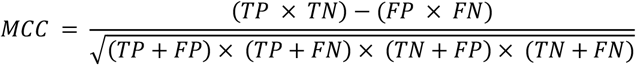

We defined the rank threshold to call positives as the rank when the MCC gives the highest value. Finally, the scoring method with the largest MCC value was adopted to define the PPI scores and detected PPIs for all subsequent analyses.

### Crystal structure analysis

The crystal structure data of the yeast 26S proteasome (57) (PDB: 6J2X) was used to calculate solvent free energy between each subunit using PDBePISA (49). The solvent free energy values were summed when multiple protein chains were available for the subunits. Kendall rank correlation was used for the statistical test. Distances from the C/N-terminal ends of the subunits were computed using the get_distance function of Pymol version 2.5.0 (Schrödinger, Inc.). The closest residue to the terminal ends available on the crystal structure was used. We adopted the closest values among subunits by considering only the α and β rings closer to the lid particle. Pearson correlation was used to compute the coefficient.

### Comparison analysis between previous DHFR-PCA datasets

Comparison analysis of the detected PPIs were carried out against previous genomic integration based DHFR-PCA (9, 58). For the protein pairs present in both BFG-PCA dataset and Tarassov *et al*, the best performing scoring method and average score of replicates were extracted from each dataset. Protein expression analysis was performed using protein abundance data (59). For each protein pair we considered the lowest expression of the pair because it is likely the limiting partner in complex formation. Mann-Whitney U-test was used for statistical tests.

### Resource availability

The DHFR-PCA plasmids for both C-terminus fusion and N-terminus fusion are available at Addgene (#177795, #177796, #177797, and #177798). The barcoded BFG-Y2H and BFG-PCA destination plasmids are available upon request.

## RESULTS

### Adapting DHFR-PCA for plasmid-based PPI detection

Gateway cloning compatible destination vectors and yeast strains were generated for BFG-PCA and are available through Addgene. We constructed a collection of 1,483 centromeric Gateway compatible plasmids with unique barcodes (893 for DHFR F[1,2] and 590 for DHFR F[3]), enabling assays of up to 526,870 protein pairs in pools using barcode fusion and sequencing (**Supplementary Table S2**). The functionality of these plasmids for DHFR-PCA and strains was first examined by performing a growth assay on 8 protein pairs consisting of 4 known PPI and 4 pairs not reported to interact. As expected, all pairs showed similar growth on the non-selective (-methotrexate) condition and pairs with reported interactions showed growth on the selective (+methotrexate) condition (**Figure 2A**). We also examined if the barcodes themselves could influence the results. We performed a separate assay on 5 protein pairs to examine if the signal is affected by the presence of different DNA barcodes. We did not observe any observable effect within the tested space **(Supplementary Figure S2**).

**Figure 2.**
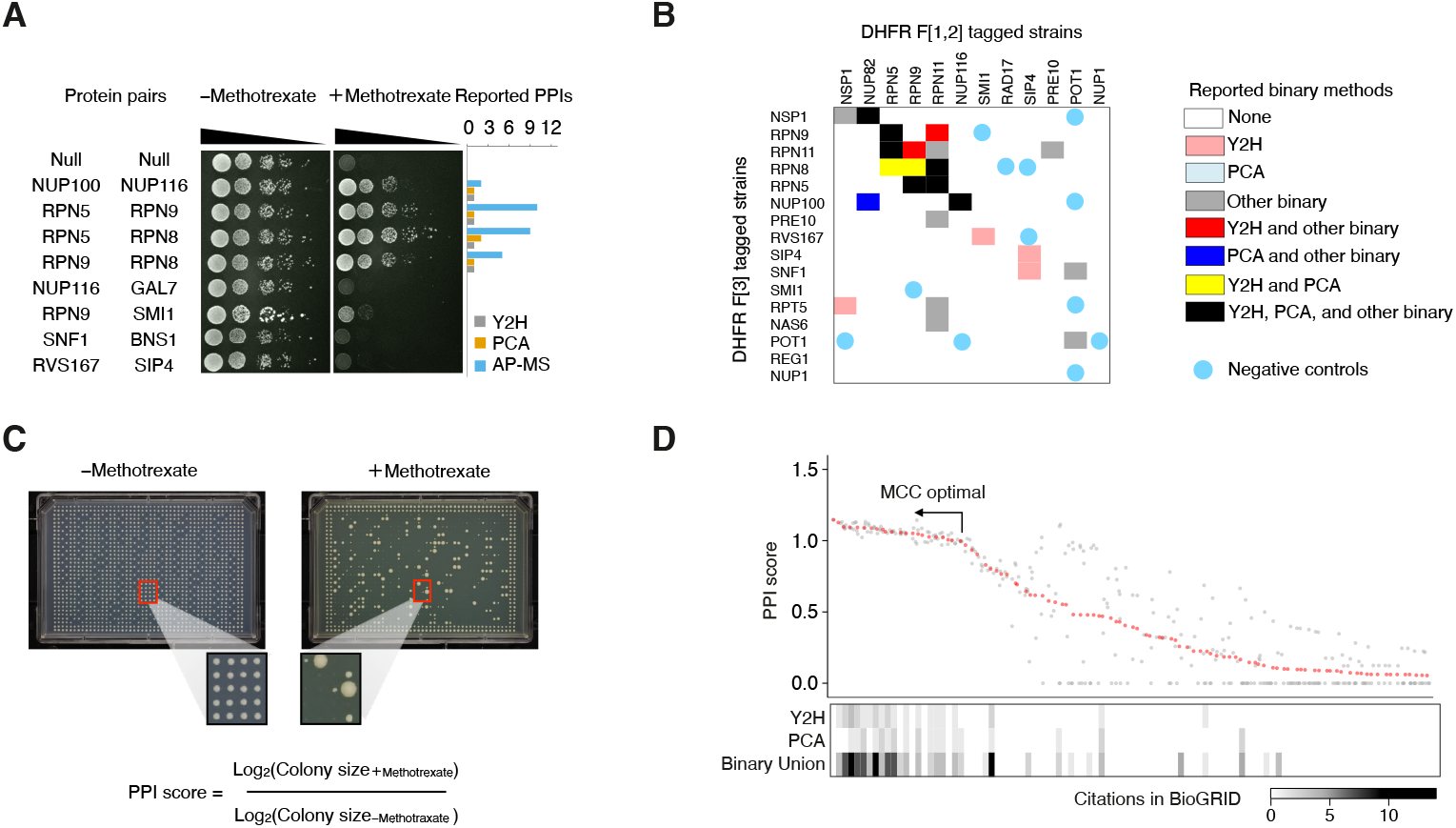
Plasmid-based DHFR-PCA captures known PPIs. (**A**) Plasmid based DHFR-PCA spot assays were performed on 8 protein pairs. Null-Null represents empty vector (destination plasmid) control. Reported PPIs in BioG RID a re shown in the barplot. While all pairs grow under no methotrexate control (-Methotrexate), only protein pairs expected to have interactions show growth with presence of methotrexate (+Methotrexate). (**B**) A subset of the query space to demonstrate the quality of plasmid-based DHFR-PCA, which was also tested by BFG-PCA (see following section). Previously known interactions are indicated in colors according to the method. (**C**) An example of a DHFR-PCA high density plate. The colony formed from replicating the same cell sample is grown on control (-Methotrexate) and selection (+Methotrexate) plates. Colony size is measured based on plate images, log-transformed and used to calculate PPI scores by fold-change between selection and control. (**D**) Result of an assay on 300 protein pairs ordered by PPI score rank. Only the top 100 protein pairs are shown. Gray dots represent replicates, and the red dot represents the 50^th^ percentile threshold used to call the ranks. The heatmap shows previously reported interactions in the BioGRID database. Binary union consists of interactions reported by Y2H, PCA, Biochemical activity, Affinity Capture-Luminescence, Reconstituted Complex, Co-crystal Structure, and FRET.

To further examine the performance of plasmid based DHFR-PCA, we performed a DHFR-PCA assay on 300 protein pairs using the established protocol on solid media (46) (**Figure 2B, C, and D**) with these plasmids. The selected space included DHFR-PCA expected positives and likely negatives for quality assessment (**Figure 2B**). Likely negative pairs were selected based on the BioGRID database (version 3.4.157) (40), with criteria including 1) no reported physical or genetic interaction for the yeast proteins or their orthologs in *Schizosaccharomyces pombe* and *Homo sapiens*, 2) no shared gene ontology terms, and 3) a distance greater than 2 edges in the PPI network. The PPI score of each pair was calculated based on colony sizes estimated from plate images (**Figure 2C**), and sorted to examine agreement with known PPIs (**Figure 2D**). Protein pairs with reported interactions were enriched for high PPI score pairs. We evaluated this by Mathew’s Correlation Coefficient (MCC), giving a value of 0.462, comparable to reported PPIs in BioGRID with either Y2H (MCC = 0.488) or PCA (MCC = 0.403). The raw PPI scores are shown in **Supplementary TableS4.**

**Figure 3.**
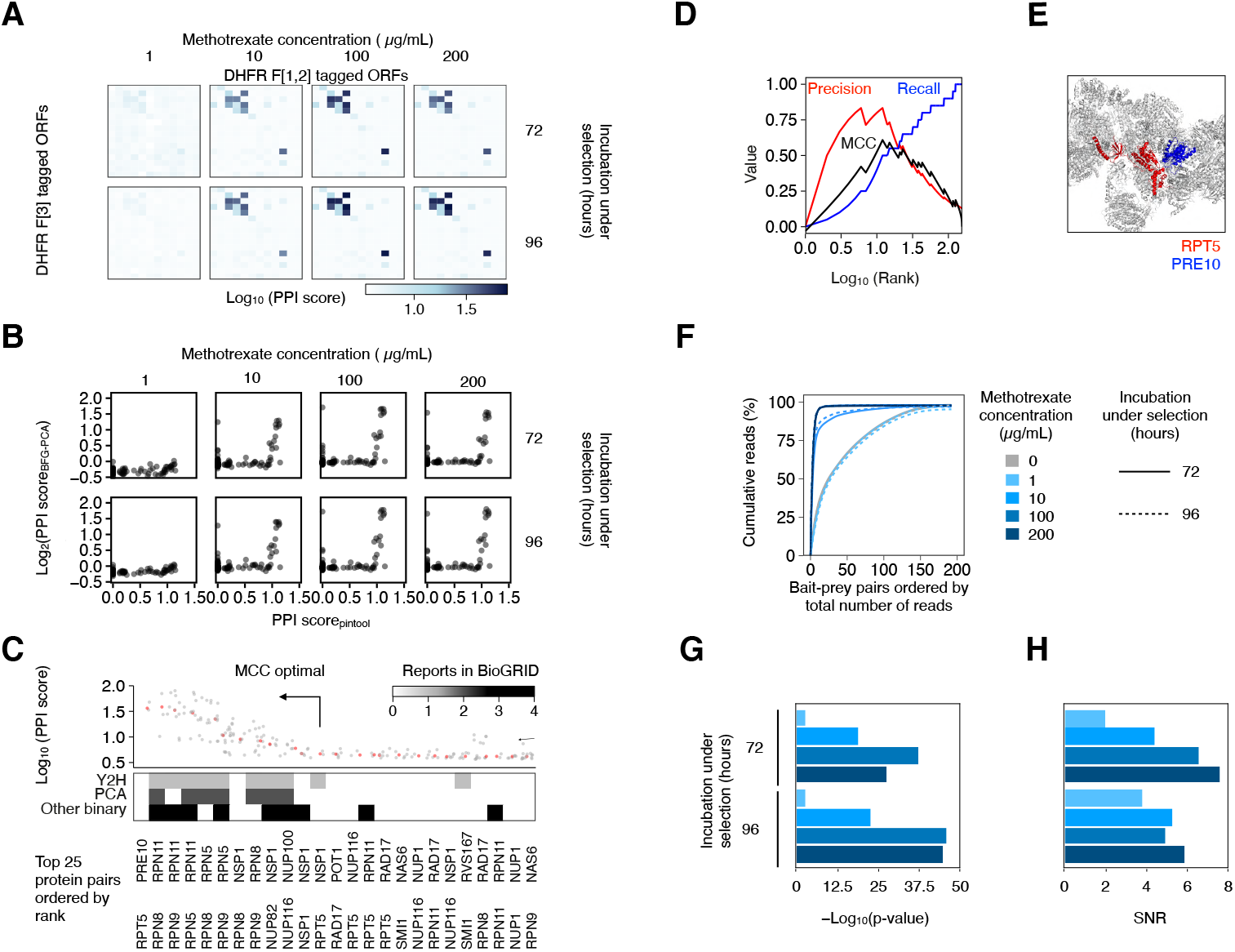
BFG-PCA screening quality is on par with one-by-one assay. (**A**) Heatmap of the BFG-PCA PPI score for each of the selection conditions. Baits and preys are ordered as in the query matrix shown in Figure 2B. (**B**) PPI scores obtained from one-by-one DHFR-PCA high density plate assay and BFG-PCA. (**C**) PPI scores ordered by rank. Methotrexate concentration of 10 μg/mL with 72 h selection is shown. Gray dots represent replicates, and the red dots represent the 50^th^ quantile of replicates used to call the rank. The heatmap represents reported interactions in the BioGRID database for Y2H, PCA, and other binary PPI detection methods (Biochemical activity, Affinity Capture-Luminescence, Reconstituted Complex, Co-crystal Structure, and FRET). (**D**) Precision/recall curve of the BFG-PCA data with methotrexate concentration of 10 μg/mL with 72 h of selection. (**E**) The top rank Pre10 and Rpt5, which had no previous binary interaction reported, are highlighted on the crystal structure of the yeast 26S proteasome (PDB: 6J2X). (**F**) Cumulative plot of raw barcode counts per protein pair under each selection condition, showing the number of protein pairs represented after sequencing. (**G & H**) The Kendall rank correlation coefficient (G), and signal to noise ratio (H) for each BFG-PCA condition against one-by-one DHFR-PCA. To compute the signal to noise ratio, the PPI scores of 12 negative control pairs and the top 10 ranked scores were averaged and used as background and signal, respectively.

### BFG-PCA screening condition optimization on 192 protein pairs

We performed a proof-of-concept BFG-PCA screening on a subset of 192 protein pairs assayed by plasmid based DHFR-PCA (**Figure 2B, 3, and Supplementary Table S4**), with the exception of 3 DHFR F[1,2] and 4 DHFR F[3] tagged constructsthat were insufficiently barcoded for BFG screening. Previous DHFR-PCA conditions used a methotrexate concentration of 200 μg/mL (9). However, Yachie et al (32) have shown that BFG-Y2H performs better when selecting under conditions less stringent than those of standard Y2H. Therefore, four concentrations (200, 100, 10 and 1 μg/mL) of methotrexate were tested to examine the optimal concentration for BFG-PCA (see **Figure 1D** for selection step). As expected, higher concentrations resulted in fewer and smaller colonies (**Supplementary Figure S3A**). Deep sequencing confirmed that the signal-to-noise ratio (SNR) increases with increasing concentration (**Supplementary Figure S3B and Figure 3A**). We compared the standard DHFR-PCA (based on colony growth) with the BFG-PCA scores (computed based on fused barcode counts). As expected, there was no relationship between the colony-based signal and BFG-PCA signal for non-interacting pairs (low colony-based signal) but a strong one above a given threshold, which corresponds to expected positive PPIs. This led to an overall positive rank correlation for all tested BFG-PCA selection conditions (Kendall rank coefficient: 1μg/mL, 72 h= 0.141; 96 h = 0.211; 10μg/mL, 72 h= 0.228; 96 h = 0.252; 100μg/mL, 72 h = 0.287; 96 h = 0.243; 200μg/mL, 72 h= 0.309; 96 h = 0.270; p<0.01, **Figure 3B and G**). The increasing correlation at higher concentrations of methotrexate and longer incubation periods contributed to higher SNRs (**Figure 3F and H**).

Among the BFG-PCA conditions tested, 10 μg/mL of methotrexate and 72 hours of selection yielded the best agreement with reported binary PPIs (**Figure 3C and D**), with a MCC of 0.61. One exception is an interaction between Pre10 and Rpt5 within the 26S proteasome complex, which had not been reported previously by any binary PPI detection method. The two proteins are neighboring within the complex when mapped to the crystal structure (**Figure 3E**), suggesting this is a true positive interaction that has been missed from previous experiments. These conditions therefore appear to be optimal among the ones tested for BFG-PCA screenings.

### BFG-PCA and BFG-Y2H screening on the Proteasome and Nuclear pore complex related proteins

Since BFG has only been implemented for Y2H, and Y2H is the most frequently used method for binary PPI screening, we compared BFG-PCA and BFG-Y2H side-by-side. We examined a space consisting of 120 proteins (34 human proteins as Y2H positive controls previously used in BFG-Y2H, 16 *S. cerevisiae* proteins used for the first demonstration of BFG-PCA, and 80 *S. cerevisiae* proteins associated with the proteasome complex and/or nuclear pore complex) with 2 barcode replicates. It is known that Y2H is less performant than DHFR-PCA to detect binary interactions within multi-subunit complexes. However, we selected this reference set as other reference sets may be less appropriate for DHFR-PCA. We performed two screenings for both BFG-PCA and BFG-Y2H (**Figure 1D**), each covering 11,232 and 10,545 bait-prey pairs, respectively. The number of barcode replicates per ORF detected in each screening were mostly 2, with some having only 1 due to loss during the cloning process (**Supplementary Figure S4A**). The distribution of bait/prey barcode abundance in the non-selective conditions, representing relative abundance of haploid strains before mating, followed a log-normal distribution in each screening as expected (**Supplementary Figure S4B**). Similarly, the relative abundance of bait-prey barcodes, representing diploid strains, followed a log-normal distribution of barcodes in the non-selective conditions (**Supplementary Figure S5**).

We computed the enrichment score ‘*s*’ as growth enrichment of each bait-prey barcode in selective conditions compared to non-selective conditions. Under selection for both BFG-PCA and BFG-Y2H, some of the bait/prey barcodes exhibited strong background noise (**Supplementary Figure S6**, **Supplementary Figure S7**, and **Supplementary Table S5**). This is a commonly known phenomenon for DB strains in Y2H where some proteins directly recruit the transcription machinery without the presence of an interaction partner (44) (**Figure 1B**). While BFG-Y2H involves a normalization for the auto-activity of problematic baits, we observed that several bait ORFs occupied >68% of all reads sequenced from selective condition libraries, which is wasted sequencing effort. Therefore, we performed an additional screening in duplicate with these strains removed for better assessment between BFG-PCA and BFG-Y2H. After removal of BFG-Y2H auto-activators, BFG-PCA and BFG-Y2H each covered 11,232 and 9,546 bait-prey pairs, respectively. When examining BFG-PCA signal data, we observed a similar but less intense auto-activity background (**Supplementary Figure S7**) which we systematically normalized when computing PPI scores. This is a known phenomenon, where some proteins interact with the DHFR fragment or the linker alone, contributing to systematic background noise (9). For implementation of either BFG-PCA or BFG-Y2H examining new baits and preys, it may therefore be necessary to first screen for a tendency for non-specific complementation or auto-activity.

The enrichment signals within replicates, both internal and screening replicates, correlated strongly in each method (**Supplementary Figure S8**), demonstrating their reproducibility. For each pair, we performed normalization of the enrichment scores based on auto-activity backgrounds (**Supplementary Figure S9A,**see methods) to obtain PPI scores. For each of the 2 screening replicates performed, each protein pair had multiple levels of internal replicates corresponding to tagging orientation, barcoded strains, and barcode fusions (**Supplementary Figure S8A**). When combining all the screening replicates and internal replicates, the average number of replicates for each protein pair was 23.8 and 21.6 for BFG-PCA and BFG-Y2H, respectively. Both screenings had over 99% of the protein pairs with more than 8 replicates (**Supplementary Figure S8F**). To call positives, we examined the best scoring method by testing average, median, and various percentile thresholds amongst the normalized score of replicates and by computing the best agreement against reported binary interactions in BioGRID (**Supplementary Figure S9B**).

As a result, we detected 92 (MCC = 0.315) and 35 (MCC = 0.296) PPIs for BFG-PCA and BFG-Y2H, respectively (**Supplementary Figure S10 and Table S6**). We compared the detected interactions of BFG-PCA and BFG-Y2H in a space of 9504 bait-prey (96 x 99 ORFs) pairs for which both data was available (**Figure 4A and B**). Although the overlap between the two methods was limited, for known binary PPIs, the PPI scores are correlated (**Figure 4C**) (R =0.12, p-value = 6.4 × 10^-3^, Kendall rank correlation). We further assessed the overall performance on the Human protein and yeast protein subsets individually (**Supplementary Figure S11 and S12**). On the Human protein subset, BFG-PCA and BFG-Y2H detected 15 (MCC =0.462), and 34 (MCC = 0.619) PPIs, respectively (**Figure 4D**, **Supplementary Figure S11**). The difference between BFG-PCA and BFG-Y2H can be explained by the fact that known Y2H positive pairs have been deliberately included in the space as positive control. In addition, no PPI data is available on human proteins screened by DHFR-PCA so we had no a priori expectation for the performance of BFG-PCA on these. On the yeast protein subset, BFG-PCA and BFG-Y2H detected 80 (MCC = 0.311) and 8 (MCC = 0.166) PPIs, respectively (**Supplementary Figure S12**). We also detected 3 and 2 cross-species PPIs by BFG-PCA and BFG-Y2H, respectively.

**Figure 4.**
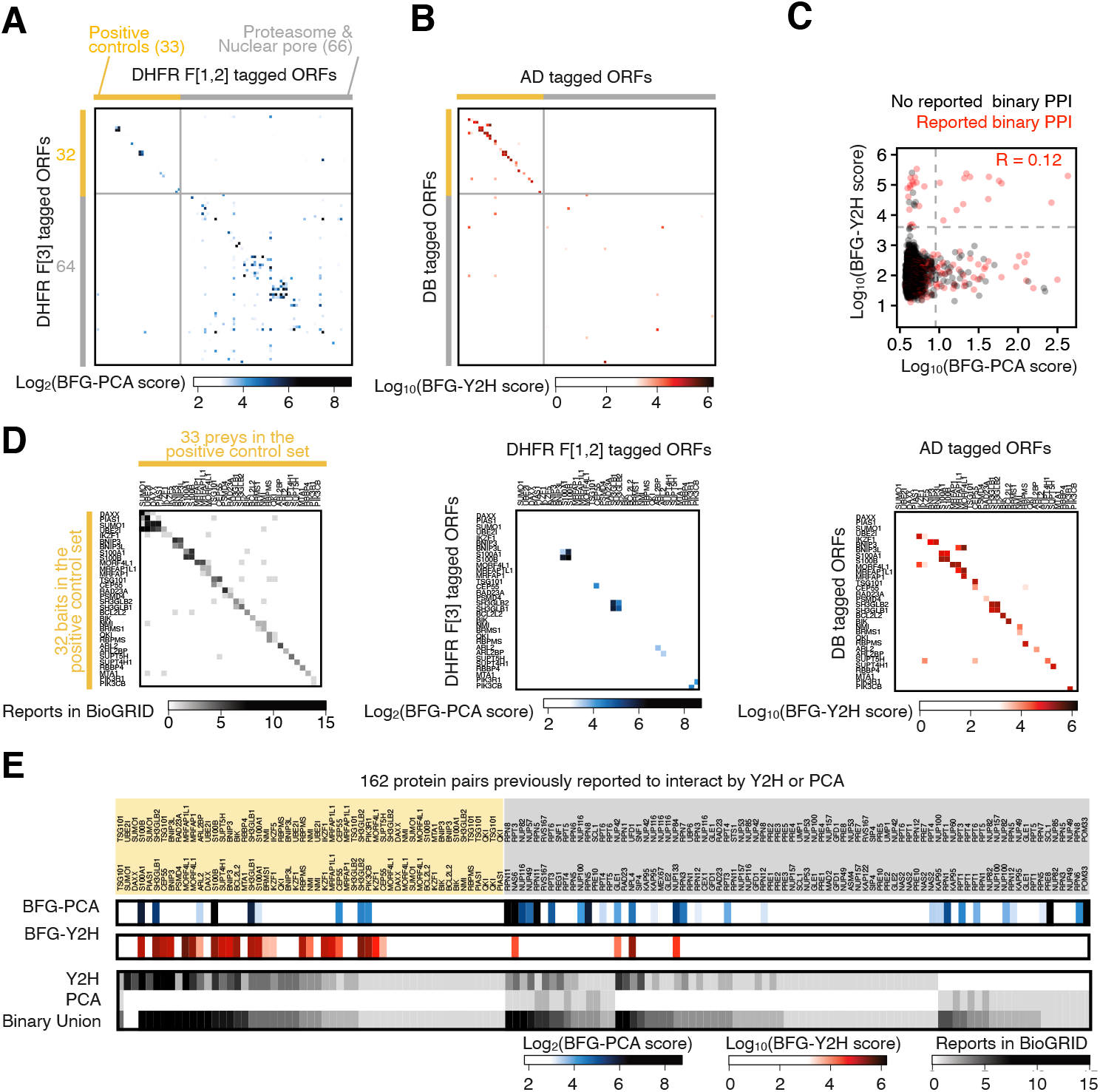
Comparative analysis of BFG-PCA and BFG-Y2H scores. (**A & B**) Heatmap representation of PPI scores for BFG-PCA (A) and BFG-Y2H (B). The ORFs present in both datasets are shown. **(C)** Scatter plot representation of BFG-PCA and BFG-Y2H scores. (**D**) Expected positive controls and subset of the data. (Left) Previously reported interactions by binary PPI detection methods. (Middle) Positive control subset of the BFG-PCA data shown in A. (Right) Positive control subset of the BFG-PCA data shown in B. (**E**) PPI scores of BFG-PCA (top) and BFG-Y2H (middle) together with reported interactions in the BioGRID database (bottom) for a subset of the screened space where the protein interaction has been reported using either Y2H or PCA.

Our reference set was biased for DHFR-PCA detectable interactions, for instance PPIs in large protein complexes. To perform a fair comparison we extracted a subset of interactions which were previously detected by either Y2H or PCA (**Figure 4E**) as those could in principle also be detected by here. BFG-PCA and BFG-Y2H captured 51.4% (19/37) and 2.7% (1/37) of interactions previously reported by PCA, and 24.1% (27/112) and 25.0% (28/112) of interactions previously reported by Y2H, respectively. Overall, BFG-PCA and BFG-Y2H detected 27.8% (37/133) and 21.1% (28/133) of the union of reported interactions by PCA and Y2H, respectively. This observation reveals that our assays did not detect all previously reported interactions. However, it is known that independent Y2H screenings often capture a fraction of detectable interactions, and multiple screenings are required for larger coverage (33, 60). From these comparisons, we can say that the sensitivity of the assays are comparable for Y2H positive interactions but BFG-PCA performs better for previously reported PCA interactions.

In conclusion, while BFG-Y2H had a higher overall ability for capturing the human PPIs, which was tailored as being positive controls for this method, BFG-PCA performed better when testing yeast protein pairs that are part of protein complexes for which Y2H sensitivity is limited, and had fewer issues with autoactivator proteins for this particular set of proteins.

### BFG-PCA captured binary PPIs in the 26S proteasome with high resolution

We further investigated whether the quantitative PPI score from BFG-PCA correlates with interaction strength. Because we used the same promoters for all subunits and all subunits within protein complexes tend to have similar protein abundance (balance between synthesis and degradation), we hypothesized that at least a fraction of the BFG-PCA signal would depend on the strength of the assemblies. We calculated the solvation free energy gain (ΔG) between subunits in the three-dimensional protein interface of crystal structure of the yeast 26S proteasome (57) (PDB: 6J2X). We observed a strong negative correlation between BFG-PCA PPI score and ΔG (R = −0.58, p-value = 1.44 ×10^−8^, Pearson correlation) (**Figure 5A**). Among the protein pairs with BFG-PCA PPI scores above the threshold to call interaction, 6 were unreported using binary PPI detection methods. We mapped these protein pairs on the crystal structure, and found that the interactions called by BFG-PCA are indeed neighboring subunits of the 26S proteasome (**Figure 5B**). These results suggest the potential of BFG-PCA to capture binary PPIs within protein complexes with high precision.

**Figure 5.**
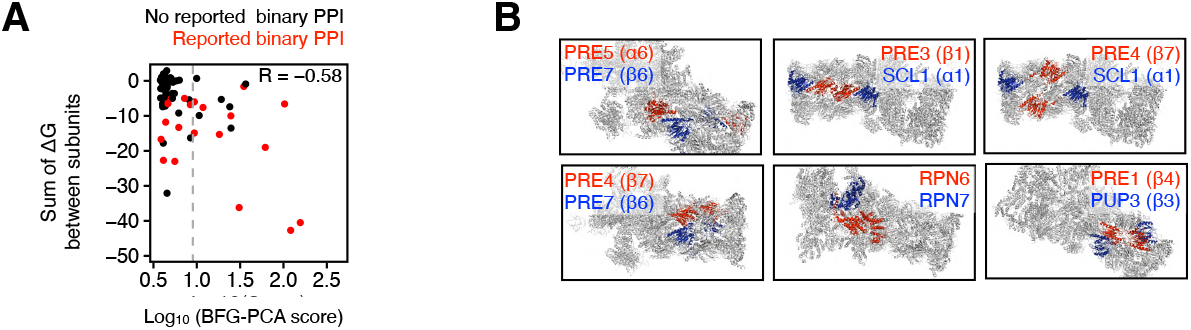
BFG-PCA PPI score agrees with ΔG between subunits within the yeast 26S proteasome. (**A**) Scatter plot of BFG-PCA score and solvation free energy gain (ΔG) upon formation of the interface between subunits. Gray dashed lines represent the threshold to call positives. Red represents PPIs in the BioGRID database reported by binary PPI detection methods. R represents Pearson correlation coefficient. (**B)** Detected positives with no previous binary PPI reports were mapped on the crystal structure (PDB:6J2X). Color for each indicated protein is shown within each image.

### BFG-PCA compared to genomic integration-based DHFR-PCA

Compared to genome-based DHFR-PCA, our plasmid constructs differ in two key components. First, the standard DHFR-PCA detects PPIs among proteins under native expression levels, whereas plasmid-based DHFR-PCA and BFG-PCA express the gene under a constitutively active *ADH1* promoter. Second, while protein-linker-DHFR fusion in previous works generally used the linker sequence (GGGS)_2_, our plasmid based linker sequence is NPAFLYKVVGGGSTS. To examine if these differences influence the detection of PPIs, we compared the interaction scores derived from DHFR-PCA with genomic integration from previous studies (9, 58) with BFG-PCA. As expected from the results reported above, BFG-PCA detected a significant number of known binary interactions which were not captured by genome-based DHFR-PCA (**Supplementary Figure S13A**). The expression levels of protein (59) with lower values within the pair (which may serve as a bottleneck for signal) was compared between each section of the scatter plot that define which PPI is detectable with each method (**Supplementary Figure S13B**). The results showed that PPI negative pairs have significantly lower expression compared to that of positive pairs, which agrees with previous literature (61). Although no significant difference in protein expression was observed between BFG-PCA specific positives and Tarassov et al.’s positives, we noticed a case of lowly expressed proteins whose expression is detectable only by BFG-PCA, Gle1, interacting with Nup42. Gle1 and Nup42 are both subunits of the nuclear pore complex, and their PPI has been reported to interact by multiple methods in both low (62–65) and high-throughput (19), but has not been detected by genome-based DHFR-PCA. The higher expression level or the modified linker or both may allow a better detection of this PPI.

We investigated whether variation in linker length or composition of polar amino acids affected the detected PPIs, we compared the BFG-PCA PPI scores to a previous effort of extending the linker sequence to (GGGS)_4_ in genome-based DHFR-PCA (58), **Supplementary Figure S14A**). Out of the four combinations of linkers they have tested, we observed that the scores obtained by standard (GGGS)_2_ for both DHFR F[1,2] and DHFR F[3] had the best agreement with BFG-PCA positives (MCC =0.50), where having an extended linker for either DHFR F[1,2] and DHFR F[3] slightly decreases the agreement (MCC = 0.48 and 0.47), and having both linkers extended drastically decreases the agreement (MCC = −0.08, **Supplementary Figure S14B).**

### Tagging orientation of DHFR fragments modifies the space of detectable PPIs

It was previously reported that the DHFR fragment position of fusion protein influences detection ability (39, 66). In the context of the proteasome for instance, we indeed observed a stronger negative correlation between BFG-PCA PPI scores and the distance between pairs of C-termini (R = −0.41, p-value = 2.04 ×10^−14^, Pearson correlation) than the N-termini (R = −0.22, p-value = 6.46 ×10^−5^, Pearson correlation) (**Figure 6A and B**). We therefore also constructed BFG-PCA plasmids amenable for DHFR N-terminal tagging. We investigated if the number of detected PPIs can be increased by using a N-terminus fusion version of DHFR-PCA. We tested interactions for 41 bait-prey pairs by spot assay with the N-terminus fusion version of plasmid-based DHFR-PCA. As a result, the N-terminus DHFR-PCA captured 7 interactions which the C-terminus tagging BFG-PCA could not capture (**Figure 6C**). Since 5 out of the 7 were detected by BFG-Y2H (N-terminus tagging), these results suggest that part of the difference between PCA and Y2H comes from tagging orientation. Another captured interaction (Rpt5p and Rpt4p) had a BFG-PCA score slightly higher compared to other tested pairs but still below the threshold to call as positives. Since the distance from C/N-terminus of this pair was 47.841 Å and 9.692 Å on the crystal structure, we suspect that the DHFR reporter reconstitution needed for cell growth was not sufficient for C-terminus DHFR fusions, but adequate for N-terminus, showing its potential for further discoveries of binary PPIs.

**Figure 6.**
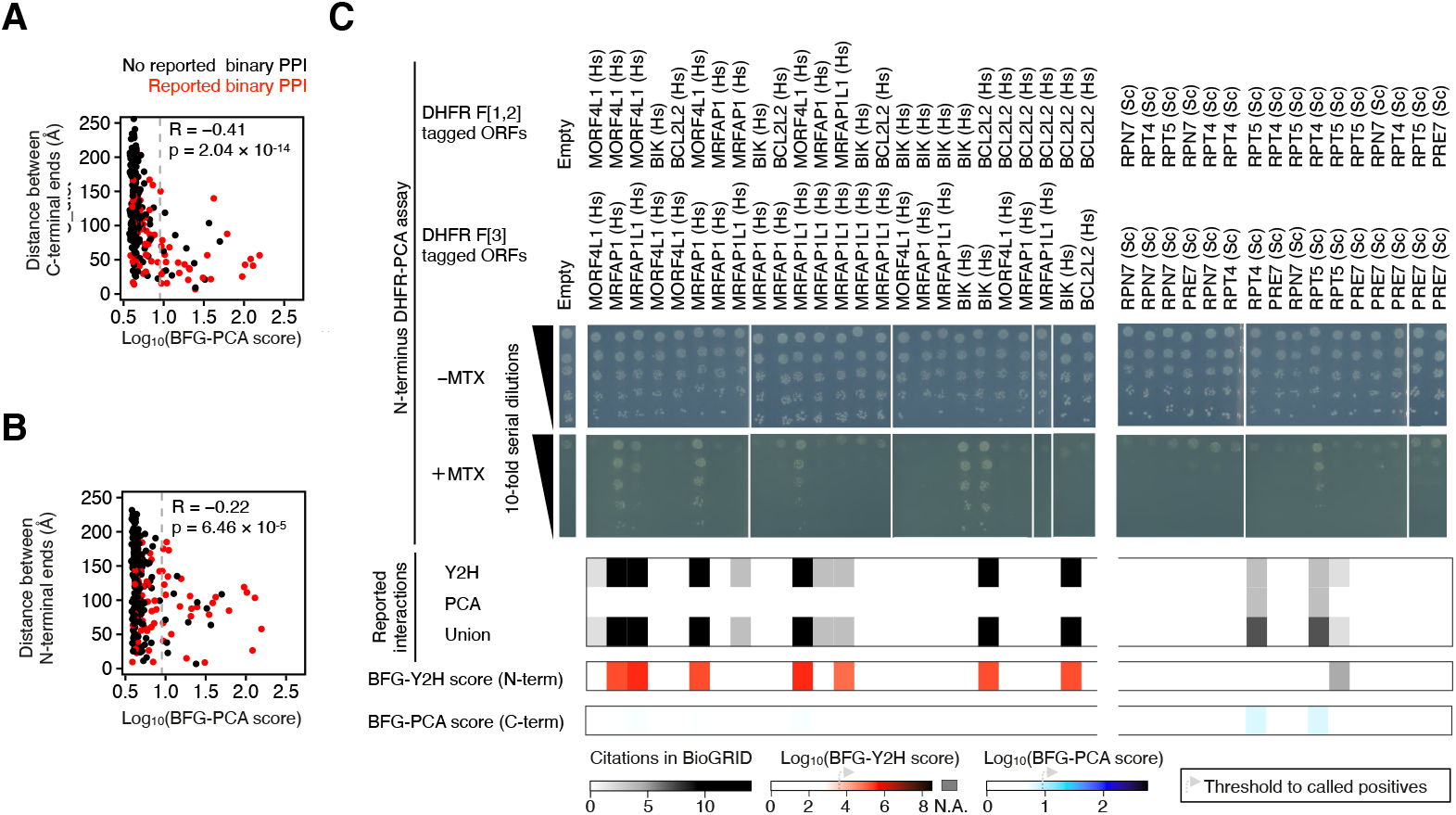
N-terminal DHFR-PCA detects BFG-Y2H specific positives. (**A & B**) Scatter plot representation of BFG-PCA PPI score and distance between the most C-terminal (A) and N-terminal (B) residue of the subunits annotated within the Yeast 26S proteasome (PDB:6J2X). Red represents PPIs in the BioGRID database reported by binary PPI detection methods. Gray dashed lines represent the threshold to call positives. R represents Pearson correlation. (**C**) Plasmid based N-terminus DHFR-PCA spot assay results for a subset of interactions screened in BFG-PCA and BFG-Y2H. Two matrices each having 5 Human proteins, and 4 Yeast proteins were tested. (Middle) Plate images from the spot assay. -MTX: Cell viability control without methotrexate. +MTX: Selection condition for PPI with 200 μg/mL methotrexate. Serial 10-fold dilution of cells starting at OD_600_nm = 0.5 was plated. (Bottom) Heatmap representation of reported binary PPIs in the BioGRID database, BFG-Y2H score, and BFG-PCA score. Data for Rpt5-Rpt5 was not available for BFG-Y2H.

## DISCUSSION

We developed a toolkit for plasmid based DHFR-PCA that exploits DNA barcode technologies for pooled screening (BFG-PCA). These tools are ready for systematic binary PPI mapping. We demonstrated the significance of BFG-PCA by screening >11,000 bait-prey pairs corresponding to 6,575 unique putative PPIs. We also performed a side-by-side comparison with BFG-Y2H for quality assessment of the method. Although it has been known that PPIs detected by DHFR-PCA and Y2H have very little overlap, no systematic comparison of the two methods has been done using the same expression promoters and the same analytical pipeline. Here, we showed that BFG-PCA and BFG-Y2H detect distinct sets of PPIs expressed from the same vector and promoter, confirming their complementarity for binary mapping. We note that BFG-PCA is significantly better at detecting yeast proteasome and nuclear pore complex related PPIs. Many reasons could explain these differences, for instance the localization of the fusion proteins. Y2H domain fused proteins are localized in the nucleus and need to have access to the chromatin and DNA to activate the expression of the selection marker, which may not always be possible for Proteasome and Nuclear Pore subunits. Further investigations would be required to characterize protein interactions detectable by BFG-PCA on a larger and more diverse set of proteins.

Previous reports described DHFR-PCA as being able to rescue the growth of cells by having as little as 25 reconstituted complexes per cell (67). Since low gene expression of an interacting partner can limit the number of DHFR reporter reconstitution, plasmid-based DHFR-PCA can in theory be more sensitive to such protein pairs than genome integration based methods. However, we found no strong evidence of such increased sensitivity of BFG-PCA compared to previous genome-based DHFR-PCA datasets in the tested protein interaction space, with a few exceptions. However, we should take into account the targeted space (the proteasome and nuclear pore complexes) in this study. It is known for instance that the subunits in the proteasome are regulated at the post transcriptional level (66), which means that higher transcription levels from the *ADH1* promoter may not influence PPIs. Also, subunits of these two protein complexes may be more expressed already than many other proteins, leaving little room for signal improvement with this promoter. In order to comprehensively assess the sensitivity of BFG-PCA on low expressed proteins, further investigation will be needed.

The C-terminus fusion DHFR-PCA constructs used in the BFG-PCA screening in our work favoured protein pairs with closer C-terminal ends. By testing N-terminus fusion DHFR-PCA constructs, we have shown that we can detect PPIs which were not detected in C-terminus fusion DHFR-PCA. Previously, it has been reported that testing all possible fusion protein orientation (C-C, C-N, N-C, and N-N fusion of bait and prey) in a nano-luciferase complementation assay can capture as many PPIs as having multiple orthogonal assays (39). Both C-terminus and N-terminus plasmid based DHFR-PCA presented here can be used and one would be able to assay all 4 of the protein fusion combinations (C-C, C-N, N-C, and N-N), increasing the PPIs detected. BFG-PCA screenings for the four fusion combinations could be performed by the same BFG-PCA haploid yeast strains prepared for C-terminus and N-terminus BFG-PCA by simply mating them in desired combinations. This enables researchers to screen PPIs with the additional C-N and N-C combinations without additional cost to prepare barcoded yeast strains, which require investments in performing BFG screenings.

In summary, the newly developed plasmid-based pooled DHFR-PCA is a binary PPI detection method orthogonal to existing assays that can expand the interactome space to be targeted in yeast but also for any species for which it is possible to clone and expressed ORF in yeast.

## DATA AVAILABILITY

The high-throughput sequencing data of this study are available at the Sequence Read Archive (PRJNA754191) of the NCBI. The raw barcode counts in this study is shown in **Supplementary Table S7**. The computed PPI scores from BFG-Y2H and BFG-PCA screening is shown in **Supplementary Table S6**. All scripts used in this study are available at (https://github.com/Landrylab/Evans-Yamamoto_et_al_2021).

## SUPPLEMENTARY DATA

Supplementary Data are available at NAR online.

## ACKNOWLEDGEMENT

The authors thank members of the Landry and Yachie Laboratories for continuous discussion. We also thank Dr. Ugo Dionne for critical reading of the manuscript.

## REFERENCES

1. Alberts, B. (1998) The cell as a collection overview of protein machines: Preparing the next generation of molecular biologists. Cell, 92, 291–294.

2. Vidal, M., Cusick, M.E. and Barabási, A.-L. (2011) Interactome Networks and Human Disease. Cell, 144, 986–998.

3. Rigaut, G., Shevchenko, A., Rutz, B., Wilm, M., Mann, M. and Séraphin, B. (1999) A generic protein purification method for protein complex characterization and proteome exploration. Nat. Biotechnol., 17, 1030–1032.

4. Ren, L., Emery, D., Kaboord, B., Chang, E. and Walid Qoronfleh, M. (2003) Improved immunomatrix methods to detect protein:protein interactions. Journal of Biochemical and Biophysical Methods, 57, 143–157.

5. Roux, K.J., Kim, D.I., Raida, M. and Burke, B. (2012) A promiscuous biotin ligase fusion protein identifies proximal and interacting proteins in mammalian cells. J. Cell Biol., 196, 801–810.

6. Gillet, L.C., Navarro, P., Tate, S., Röst, H., Selevsek, N., Reiter, L., Bonner, R. and Aebersold, R. (2012) Targeted data extraction of the MS/MS spectra generated by data-independent acquisition: a new concept for consistent and accurate proteome analysis. Mol. Cell. Proteomics, 11, O111.016717.

7. Fields, S. and Song, O. (1989) A novel genetic system to detect protein-protein interactions. Nature, 340, 245–246.

8. Hu, C.-D., Chinenov, Y. and Kerppola, T.K. (2002) Visualization of interactions among bZIP and Rel family proteins in living cells using bimolecular fluorescence complementation. Mol. Cell, 9, 789–798.

9. Tarassov, K., Messier, V., Landry, C.R., Radinovic, S., Serna Molina, M.M., Shames, I., Malitskaya, Y., Vogel, J., Bussey, H. and Michnick, S.W. (2008) An in vivo map of the yeast protein interactome. Science, 320, 1465–1470.

10. Rhee, H.-W., Zou, P., Udeshi, N.D., Martell, J.D., Mootha, V.K., Carr, S.A. and Ting, A.Y. (2013) Proteomic mapping of mitochondria in living cells via spatially restricted enzymatic tagging. Science, 339, 1328–1331.

11. Go, C.D., Knight, J.D.R., Rajasekharan, A., Rathod, B., Hesketh, G.G., Abe, K.T., Youn, J.-Y., Samavarchi-Tehrani, P., Zhang, H., Zhu, L.Y., et al. (2021) A proximity-dependent biotinylation map of a human cell. Nature, 595, 120–124.

12. Kristensen, A.R., Gsponer, J. and Foster, L.J. (2012) A high-throughput approach for measuring temporal changes in the interactome. Nat. Methods, 9, 907–909.

13. Salas, D., Stacey, R.G., Akinlaja, M. and Foster, L.J. (2020) Next-generation Interactomics: Considerations for the Use of Co-elution to Measure Protein Interaction Networks. Mol. Cell. Proteomics, 19, 1–10.

14. Yu, C. and Huang, L. (2018) Cross-Linking Mass Spectrometry: An Emerging Technology for Interactomics and Structural Biology. Anal. Chem., 90, 144–165.

15. Bruce, J.E. (2012) In vivo protein complex topologies: sights through a cross-linking lens. Proteomics, 12, 1565–1575.

16. Venkatesan, K., Rual, J.-F., Vazquez, A., Stelzl, U., Lemmens, I., Hirozane-Kishikawa, T., Hao, T., Zenkner, M., Xin, X., Goh, K.-I., et al. (2009) An empirical framework for binary interactome mapping. Nat. Methods, 6, 83–90.

17. Luck, K., Kim, D.-K., Lambourne, L., Spirohn, K., Begg, B.E., Bian, W., Brignall, R., Cafarelli, T., Campos-Laborie, F.J., Charloteaux, B., et al. (2020) A reference map of the human binary protein interactome. Nature, 580, 402–408.

18. Rolland, T., Taşan, M., Charloteaux, B., Pevzner, S.J., Zhong, Q., Sahni, N., Yi, S., Lemmens, I., Fontanillo, C., Mosca, R., et al. (2014) A proteome-scale map of the human interactome network. Cell, 159, 1212–1226.

19. Yu, H., Braun, P., Yildirim, M.A., Lemmens, I., Venkatesan, K., Sahalie, J., Hirozane-Kishikawa, T., Gebreab, F., Li, N., Simonis, N., et al. (2008) High-quality binary protein interaction map of the yeast interactome network. Science, 322, 104–110.

20. Li, S. (2004) A Map of the Interactome Network of the Metazoan C. elegans. Science, 303, 540–543.

21. Rajagopala, S.V., Sikorski, P., Kumar, A., Mosca, R., Vlasblom, J., Arnold, R., Franca-Koh, J., Pakala, S.B., Phanse, S., Ceol, A., et al. (2014) The binary protein-protein interaction landscape of Escherichia coli. Nat. Biotechnol., 32, 285–290.

22. Arabidopsis Interactome Mapping Consortium (2011) Evidence for network evolution in an Arabidopsis interactome map. Science, 333, 601–607.

23. Giot, L., Bader, J.S., Brouwer, C., Chaudhuri, A., Kuang, B., Li, Y., Hao, Y.L., Ooi, C.E., Godwin, B., Vitols, E., et al. (2003) A protein interaction map of Drosophila melanogaster. Science, 302, 1727–1736.

24. Smith, L.M., The Consortium for Top Down Proteomics and Kelleher, N.L. (2013) Proteoform: a single term describing protein complexity. Nature Methods, 10, 186–187.

25. Aebersold, R., Agar, J.N., Amster, I.J., Baker, M.S., Bertozzi, C.R., Boja, E.S., Costello, C.E., Cravatt, B.F., Fenselau, C., Garcia, B.A., et al. (2018) How many human proteoforms are there? Nat. Chem. Biol., 14, 206–214.

26. Sahni, N., Yi, S., Zhong, Q., Jailkhani, N., Charloteaux, B., Cusick, M.E. and Vidal, M. (2013) Edgotype: a fundamental link between genotype and phenotype. Curr. Opin. Genet. Dev., 23, 649–657.

27. Corominas, R., Yang, X., Lin, G.N., Kang, S., Shen, Y., Ghamsari, L., Broly, M., Rodriguez, M., Tam, S., Trigg, S.A., et al. (2014) Protein interaction network of alternatively spliced isoforms from brain links genetic risk factors for autism. Nat. Commun., 5, 3650.

28. Yu, H., Tardivo, L., Tam, S., Weiner, E., Gebreab, F., Fan, C., Svrzikapa, N., Hirozane-Kishikawa, T., Rietman, E., Yang, X., et al. (2011) Next-generation sequencing to generate interactome datasets. Nat. Methods, 8, 478–480.

29. Schlecht, U., Miranda, M., Suresh, S., Davis, R.W. and St Onge, R.P. (2012) Multiplex assay for condition-dependent changes in protein-protein interactions. Proc. Natl. Acad. Sci. U. S. A., 109, 9213–9218.

30. Lewis, J.D., Wan, J., Ford, R., Gong, Y., Fung, P., Nahal, H., Wang, P.W., Desveaux, D. and Guttman, D.S. (2012) Quantitative Interactor Screening with next-generation Sequencing (QIS-Seq) identifies Arabidopsis thaliana MLO2 as a target of the Pseudomonas syringae type III effector HopZ2. BMC Genomics, 13, 8.

31. Weimann, M., Grossmann, A., Woodsmith, J., Özkan, Z., Birth, P., Meierhofer, D., Benlasfer, N., Valovka, T., Timmermann, B., Wanker, E.E., et al. (2013) A Y2H-seq approach defines the human protein methyltransferase interactome. Nat. Methods, 10, 339–342.

32. Yachie, N., Petsalaki, E., Mellor, J.C., Weile, J., Jacob, Y., Verby, M., Ozturk, S.B., Li, S., Cote, A.G., Mosca, R., et al. (2016) Pooled-matrix protein interaction screens using Barcode Fusion Genetics. Mol. Syst. Biol., 12, 863.

33. Trigg, S.A., Garza, R.M., MacWilliams, A., Nery, J.R., Bartlett, A., Castanon, R., Goubil, A., Feeney, J., O’Malley, R., Huang, S.-S.C., et al. (2017) CrY2H-seq: a massively multiplexed assay for deep-coverage interactome mapping. Nat. Methods, 14, 819–825.

34. Schlecht, U., Liu, Z., Blundell, J.R., St Onge, R.P. and Levy, S.F. (2017) A scalable double-barcode sequencing platform for characterization of dynamic protein-protein interactions. Nat. Commun., 8, 15586.

35. Yang, F., Lei, Y., Zhou, M., Yao, Q., Han, Y., Wu, X., Zhong, W., Zhu, C., Xu, W., Tao, R., et al. (2018) Development and application of a recombination-based library versus library high-throughput yeast two-hybrid (RLL-Y2H) screening system. Nucleic Acids Res., 46, e17.

36. Yang, J.-S., Garriga-Canut, M., Link, N., Carolis, C., Broadbent, K., Beltran-Sastre, V., Serrano, L. and Maurer, S.P. (2018) rec-YnH enables simultaneous many-by-many detection of direct protein-protein and protein-RNA interactions. Nat. Commun., 9, 3747.

37. Liu, Z., Miller, D., Li, F., Liu, X. and Levy, S.F. (2020) A large accessory protein interactome is rewired across environments. Elife, 9.

38. Braun, P., Tasan, M., Dreze, M., Barrios-Rodiles, M., Lemmens, I., Yu, H., Sahalie, J.M., Murray, R.R., Roncari, L., de Smet, A.-S., et al. (2009) An experimentally derived confidence score for binary protein-protein interactions. Nat. Methods, 6, 91–97.

39. Choi, S.G., Olivet, J., Cassonnet, P., Vidalain, P.-O., Luck, K., Lambourne, L., Spirohn, K., Lemmens, I., Dos Santos, M., Demeret, C., et al. (2019) Maximizing binary interactome mapping with a minimal number of assays. Nat. Commun., 10, 3907.

40. Stark, C. (2006) BioGRID: a general repository for interaction datasets. Nucleic Acids Research, 34, D535–D539.

41. Buntru, A., Trepte, P., Klockmeier, K., Schnoegl, S. and Wanker, E.E. (2016) Current Approaches Toward Quantitative Mapping of the Interactome. Front. Genet., 7, 74.

42. Celaj, A., Schlecht, U., Smith, J.D., Xu, W., Suresh, S., Miranda, M., Aparicio, A.M., Proctor, M., Davis, R.W., Roth, F.P., et al. (2017) Quantitative analysis of protein interaction network dynamics in yeast. Mol. Syst. Biol., 13, 934.

43. Amberg, D.C., Burke, D. and Strathern, J.N. (2005) Methods in Yeast Genetics: A Cold Spring Harbor Laboratory Course Manual CSHL Press.

44. Gibson, D.G., Young, L., Chuang, R.-Y., Venter, J.C., Hutchison, C.A., 3rd and Smith, H.O. (2009) Enzymatic assembly of DNA molecules up to several hundred kilobases. Nat. Methods, 6, 343–345.

45. James, P., Halladay, J. and Craig, E.A. (1996) Genomic Libraries and a Host Strain Designed for Highly Efficient Two-Hybrid Selection in Yeast. Genetics, 144, 1425–1436.

46. Marchant, A., Cisneros, A.F., Dubé, A.K., Gagnon-Arsenault, I., Ascencio, D., Jain, H., Aubé, S., Eberlein, C., Evans-Yamamoto, D., Yachie, N., et al. (2019) The role of structural pleiotropy and regulatory evolution in the retention of heteromers of paralogs. Elife, 8.

47. Rual, J.-F., Venkatesan, K., Hao, T., Hirozane-Kishikawa, T., Dricot, A., Li, N., Berriz, G.F., Gibbons, F.D., Dreze, M., Ayivi-Guedehoussou, N., et al. (2005) Towards a proteome-scale map of the human protein-protein interaction network. Nature, 437, 1173–1178.

48. Altschul, S.F., Gish, W., Miller, W., Myers, E.W. and Lipman, D.J. (1990) Basic local alignment search tool. Journal of Molecular Biology, 215, 403–410.

49. Krissinel, E. and Henrick, K. (2007) Inference of macromolecular assemblies from crystalline state. J. Mol. Biol., 372, 774–797.

50. Walhout, A.J.M., Temple, G.F., Brasch, M.A., Hartley, J.L., Lorson, M.A., van den Heuvel, S. and Vidal, M. (2000) GATEWAY recombinational cloning: Application to the cloning of large numbers of open reading frames or ORFeomes. Methods in Enzymology, 10.1016/s0076-6879(00)28419-x.

51. Dreze, M., Monachello, D., Lurin, C., Cusick, M.E., Hill, D.E., Vidal, M. and Braun, P. (2010) High-Quality Binary Interactome Mapping. Methods in Enzymology, 10.1016/s0076-6879(10)70012-4.

52. Rochette, S., Diss, G., Filteau, M., Leducq, J.-B., Dubé, A.K. and Landry, C.R. (2015) Genome-wide protein-protein interaction screening by protein-fragment complementation assay (PCA) in living cells. J. Vis. Exp., 10.3791/52255.

53. Lõoke, M., Kristjuhan, K. and Kristjuhan, A. (2011) Extraction of genomic DNA from yeasts for PCR-based applications. Biotechniques, 50, 325–328.

54. The UniProt Consortium (2017) UniProt: the universal protein knowledgebase. Nucleic Acids Res., 45, D158–D169.

55. Gelperin, D.M., White, M.A., Wilkinson, M.L., Kon, Y., Kung, L.A., Wise, K.J., Lopez-Hoyo, N., Jiang, L., Piccirillo, S., Yu, H., et al. (2005) Biochemical and genetic analysis of the yeast proteome with a movable ORF collection. Genes Dev., 19, 2816–2826.

56. Matthews, B.W. (1975) Comparison of the predicted and observed secondary structure of T4 phage lysozyme. Biochim. Biophys. Acta, 405, 442–451.

57. Ding, Z., Xu, C., Sahu, I., Wang, Y., Fu, Z., Huang, M., Wong, C.C.L., Glickman, M.H. and Cong, Y. (2019) Structural Snapshots of 26S Proteasome Reveal Tetraubiquitin-Induced Conformations. Molecular Cell, 73, 1150–1161.e6.

58. Chrétien, A.-È., Gagnon-Arsenault, I., Dubé, A.K., Barbeau, X., Després, P.C., Lamothe, C., Dion-Côté, A.-M., Lagüe, P. and Landry, C.R. (2018) Extended linkers improve the detection of proteinprotein interactions (PPIs) by dihydrofolate reductase protein-fragment complementation assay (DHFR PCA) in living cells. Mol. Cell. Proteomics, 17, 549.

59. Ho, B., Baryshnikova, A. and Brown, G.W. (2018) Unification of Protein Abundance Datasets Yields a Quantitative Saccharomyces cerevisiae Proteome. Cell Syst, 6, 192–205.e3.

60. Ito, T., Chiba, T., Ozawa, R., Yoshida, M., Hattori, M. and Sakaki, Y. (2001) A comprehensive two-hybrid analysis to explore the yeast protein interactome. Proc. Natl. Acad. Sci. U. S. A., 98, 4569–4574.

61. Grigoriev, A. (2001) A relationship between gene expression and protein interactions on the proteome scale: analysis of the bacteriophage T7 and the yeast Saccharomyces cerevisiae. Nucleic Acids Res., 29, 3513–3519.

62. Murphy, R. and Wente, S.R. (1996) An RNA-export mediator with an essential nuclear export signal. Nature, 383, 357–360.

63. Strahm, Y. (1999) The RNA export factor Gle1p is located on the cytoplasmic fibrils of the NPC and physically interacts with the FG-nucleoporin Rip1p, the DEAD-box protein Rat8p/Dbp5p and a new protein Ymr255p. The EMBO Journal, 18, 5761–5777.

64. Alber, F., Dokudovskaya, S., Veenhoff, L.M., Zhang, W., Kipper, J., Devos, D., Suprapto, A., Karni-Schmidt, O., Williams, R., Chait, B.T., et al. (2007) Determining the architectures of macromolecular assemblies. Nature, 450, 683–694.

65. Adams, R.L., Mason, A.C., Glass, L., Aditi and Wente, S.R. (2018) Correction to: Nup42 and IP6 coordinate Gle1 stimulation of Dbp5/DDX19B for mRNA export in yeast and human cells. Traffic, 19, 650.

66. MacDonald, M.L., Lamerdin, J., Owens, S., Keon, B.H., Bilter, G.K., Shang, Z., Huang, Z., Yu, H., Dias, J., Minami, T., et al. (2006) Identifying off-target effects and hidden phenotypes of drugs in human cells. Nature Chemical Biology, 2, 329–337.

67. Remy, I. and Michnick, S.W. (1999) Clonal selection and in vivo quantitation of protein interactions with protein-fragment complementation assays. Proc. Natl. Acad. Sci. U. S. A., 96, 5394–5399.

