## Supplementary Notes and Figures for "Barcode Fusion Genetics-Protein-fragment Complementation Assay (BFG-PCA): tools and resources that expand the potential for binary protein interaction discovery"

### Table of contents


| | |
|---|---|
| Note S1 | Confirmation of BFG compatibility in BFG-PCA strains |
| Note S2 | Media composition used in this study |
| Figure S1 | DHFR-PCA plasmid and strains for BFG |
| Figure S2 | DNA barcodes do not affect the DHFR-PCA signal |
| Figure S3 | Increasing methotrexate concentration increases the signal-to-noise ratio |
| Figure S4 | Raw frequency distribution of bait/prey barcodes in BFG-PCA and BFG-Y2H screenings |
| Figure S5 | Raw frequency distribution of fused bait-prey barcodes in BFG-PCA and BFG-Y2H screenings |
| Figure S6 | Enrichment of bait-prey barcodes in selective conditions compared to their respective non-selection conditions. |
| Figure S7 | Comparison of auto-activity on BFG-Y2H and BFG-PCA |
| Figure S8 | Correlation of enrichment score between replicates |
| Figure S9 | Optimal MCC on normalization and scoring methods |
| Figure S10 | Quality of BFG-PCA and BFG-Y2H screenings |
| Figure S11 | Detected interactions in the Human protein subset |
| Figure S12 | Detected interactions in the Yeast protein subset |
| Figure S13 | BFG-PCA scores compared to previous DHFR-PCA datasets |
| Figure S14 | BFG-PCA scores compared to DHFR-PCA with extended linkers |

**Note S1. Confirmation of BFG compatibility in BFG-PCA strains.**

To confirm the BFG capability of the BFG-PCA strains YY3094 and YY3095, we induced Cre mediated barcode fusion of barcoded DHFR-PCA plasmids as demonstrated in Yachie et al. (Yachie et al. 2016), and compared the intra-cellular BFG event to Y2H and BFG-Y2H strains. While individual haploid strains or mix of lysates do not show any barcode fusion product, BFG-Y2H and BFG-PCA diploid strains with both the $tetO_2$ promoter regulated Cre cassette and rtTA necessary for doxycycline induced expression (**Figure S1A**) show amplification of fused barcode products (**Figure S1B**). This shows that the generated strains are functional for BFG-PCA screenings.

**Note S2. Media composition used in this study.**

YPAD

| Component | Quantity per L |
|---|---|
| Yeast extract | 10 g |
| Bacto peptone | 20 g |
| Glucose | 20 g |
| Adenine hemisulfate | 40 mg |
| ddH2O | Up to 1L |
| Agar (for plates) | 15 g |
| Autoclave and pour | |

SC-Trp+Ade

| Component | Quantity per L |
|---|---|
| SC amino acid dropout mix (-LWH) | 1.9 g |
| Yeast nitrogen base with ammonium sulfate | 6.7 g |
| ddH2O | 919 mL |
| NaOH | Adjust to pH 5.9 |
| Agar (for plates) | 30 g |
| Autoclave | |
| 100 mM Leucine solution | 8 mL |
| 100 mM Histidine solution | 8 mL |
| 40% (w/v) Glucose solution | 50 mL |
| 12 mg/mL Adenine solution | 15 mL |

SC-Leu+Ade

| Component | Quantity per L |
|---|---|
| SC amino acid dropout mix (-LWH) | 1.9 g |
| Yeast nitrogen base with ammonium sulfate | 6.7 g |
| ddH2O | 919 mL |
| NaOH | Adjust to pH 5.9 |
| Agar (for plates) | 30 g |
| Autoclave | |
| 100 mM Tryptophan solution | 8 mL |
| 100 mM Histidine solution | 8 mL |
| 40% (w/v) Glucose solution | 50 mL |
| 12 mg/mL Adenine solution | 15 mL |

SC-Trp-Leu+10xHis+Ade

| Component | Quantity per L |
|---|---|
| SC amino acid dropout mix (-LWH) | 1.9 g |
| Yeast nitrogen base with ammonium sulfate | 6.7 g |
| ddH2O | 847 mL |
| NaOH | Adjust to pH 5.9 |
| Agar (for plates) | 30 g |
| Autoclave | |
| 100 mM Histidine solution | 80 mL |
| 40% (w/v) Glucose solution | 50 mL |
| 12 mg/mL Adenine solution | 15 mL |

SC-Trp-Leu-His+Ade

| Component | Quantity per L |
| --- | --- |
| SC amino acid dropout mix (-LWH) | 1.9 g |
| Yeast nitrogen base with ammonium sulfate | 6.7 g |
| ddH2O | 911 mL |
| NaOH | Adjust to pH 5.9 |
| Agar | 30 g |
| Autoclave | |
| 40% (w/v) Glucose solution | 50 mL |
| 12 mg/mL Adenine solution | 15 mL |

SC-Trp-Leu-His+Ade+1mM 3AT

| Component | Quantity per L |
| --- | --- |
| SC amino acid dropout mix (-LWH) | 1.9 g |
| Yeast nitrogen base with ammonium sulfate | 6.7 g |
| ddH2O | 910 mL |
| NaOH | Adjust to pH 5.9 |
| Agar | 30 g |
| Autoclave, wait until cools down to 55˚C | |
| 40% (w/v) Glucose solution | 50 mL |
| 12 mg/mL Adenine solution | 15 mL |
| 1M 3-AT | 1000 µL |

SC-Trp-Leu-Ade+DMSO

| Component | Quantity per L |
|---|---|
| In Flask A | |
| Yeast nitrogen base without ammonium sulfate | 6.69 g |
| ddH2O | 412 mL |
| In Flask B | |
| dH2O | 412 mL |
| Noble Agar | 25 g |
| Autoclave | |
| Let stand at 55℃ for 30min | |
| Combine flask_A and flask_B in flask_B | |
| 2M Glucose solution | 56 mL |
| 10x AA DO mix solution (-AWL) | 100 mL |
| DMSO | 20 mL |

SC-Trp-Leu-Ade+methotrexate

| Component | Quantity per L |
|---|---|
| In Flask A | |
| Yeast nitrogen base without ammonium sulfate | 6.69 g |
| ddH2O | 412 mL |
| In Flask B | |
| dH2O | 412 mL |
| Noble Agar | 25 g |
| Autoclave | |
| Let stand at 55℃ for 30min | |
| Combine flask_A and flask_B in flask_B | |
| 2M Glucose solution | 56 mL |
| 10x AA DO mix solution (-AWL) | 100 mL |
| 10mg/mL MTX dissolved in DMSO | 20 mL |

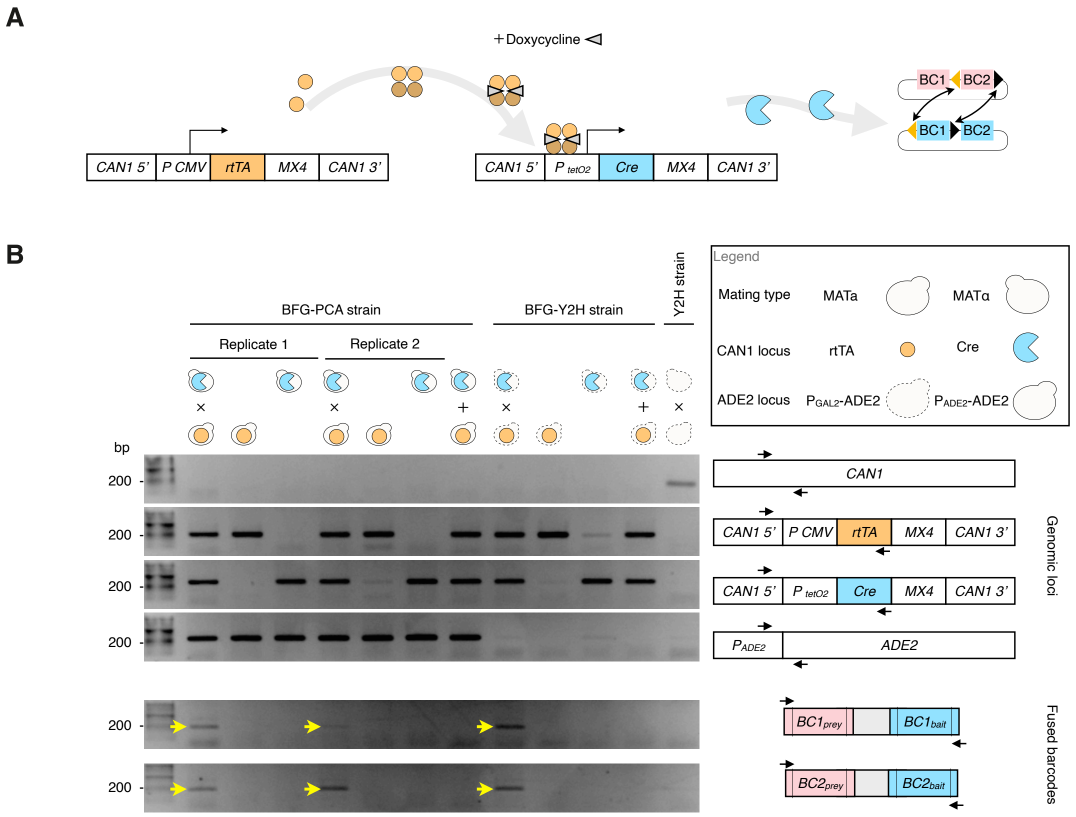


**Figure S1. DHFR-PCA plasmids and strains undergo barcode fusion upon Cre expression.**

A. Illustration of Tet-On system based Cre recombinase expression. The reverse tetracycline transactivator (rtTA) is constitutively expressed. With doxycycline, rtTA is activated, binds to the Tet-On promoter and transcribes Cre recombinase. Cre recombine LoxP and Lox2272 recognition sites for barcode fusion.
B. BFG-PCA strains (YY3094 and YY3095) were tested for barcode fusion capability under doxycycline induction. BFG-Y2H strains (RY1010 and RY1030), and Y2H strains (Y8800 and Y8930) were used as positive and negative controls, respectively. The cross between cartoons of haploids represents mated diploids. Both BFG-Y2H and BFG-PCA diploids have rtTA and Cre cassettes after mating, where the haploids only have either cassette. Y2H strains have neither rtTA and Cre. The plus between haploids represents mixing lysates of the haploids, a control to ensure the recombination happening within the cell, and not after cell lysis. BFG-PCA strains were generated from BFG-Y2H strains, but have the wild type ADE2 allele restored. The top 4 rows of gel images genotypes the genomic elements CAN1, rtTA, Cre, and ADE2 loci. The last 2 rows of genotypes fused barcode product, BC1-BC1 fusion product and BC2-BC2 fusion product. For each row, the schematic of primer annealing is illustrated on the right. Detected fused barcode products are marked with a yellow arrow.

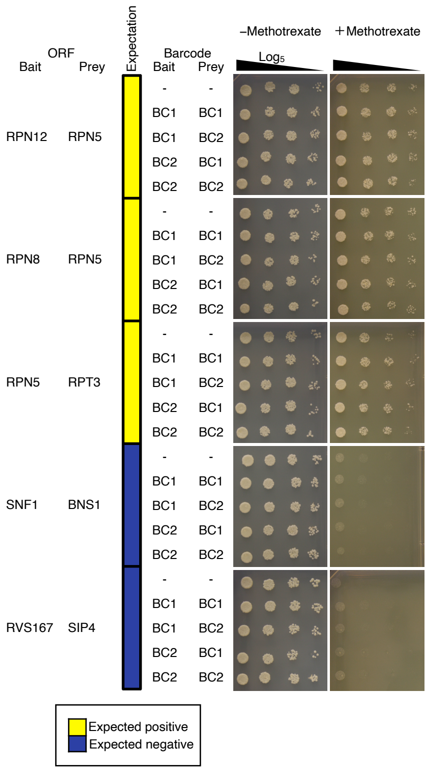


**Figure S2. DNA barcodes do not affect the DHFR-PCA signal.**

Spot assay of DHFR-PCA with and without DNA barcodes for 5 protein pairs. Four DNA barcode combinations were tested for each pair. Cells were spotted in 5-fold dilution series starting from $OD_{600nm}$ = 1.0. Yellow: Expected positives which have detected reports using DHFR-PCA. Navy: Expected negatives with no interactions reported and having distinct GO terms.

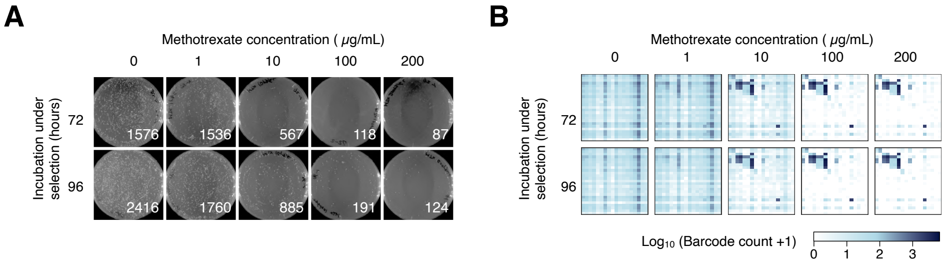


**Figure S3. Increasing methotrexate concentration increases the signal-to-noise ratio.**

A. Plate observation after BFG-PCA selection under varying methotrexate concentration and time points. Representative plates are shown with the average CFU count shown on the bottom right corner (n=2). Note that only visible colonies could be counted at each condition.

B. Heatmap representation of raw barcode counts from deep sequencing for each of the tested BFG-PCA conditions. The bait and prey barcodes are ordered as in the query matrix shown in Figure EV1B.

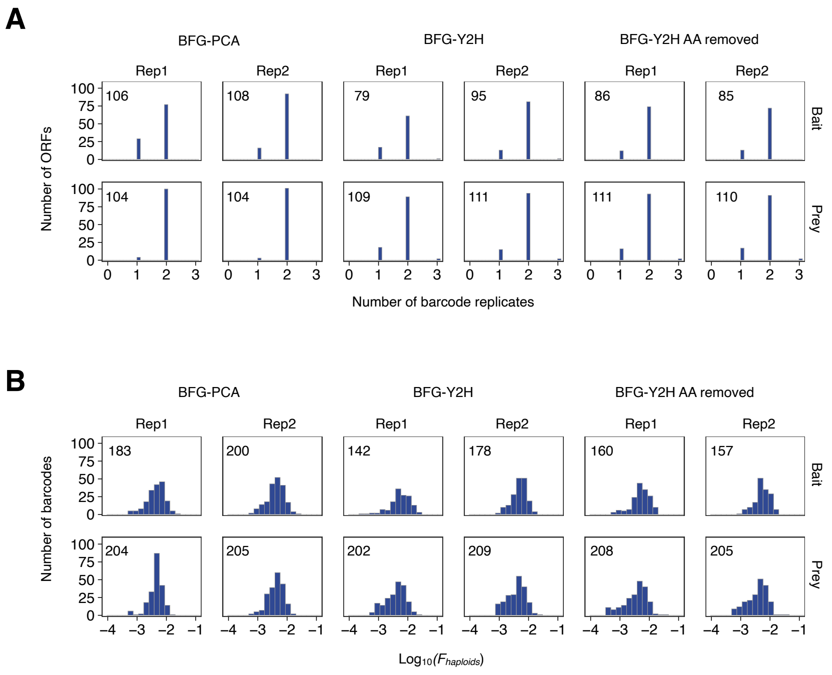


**Figure S4. Raw frequency distribution of bait/prey barcodes in BFG-PCA and BFG-Y2H screenings.**

A. Histogram representation of the number of unique barcodes assigned to each ORF. The total number of ORFs are indicated at the top left corner.
B. Histogram representation of raw abundance for each bait/prey barcode. Since the sequencing readouts are fused (bait-prey) barcodes, we aggregated the count of barcodes according to each bait/prey barcode to calculate marginal abundance of bait/prey barcodes. The total number of barcodes detected in the screening is indicated at the top right corner.

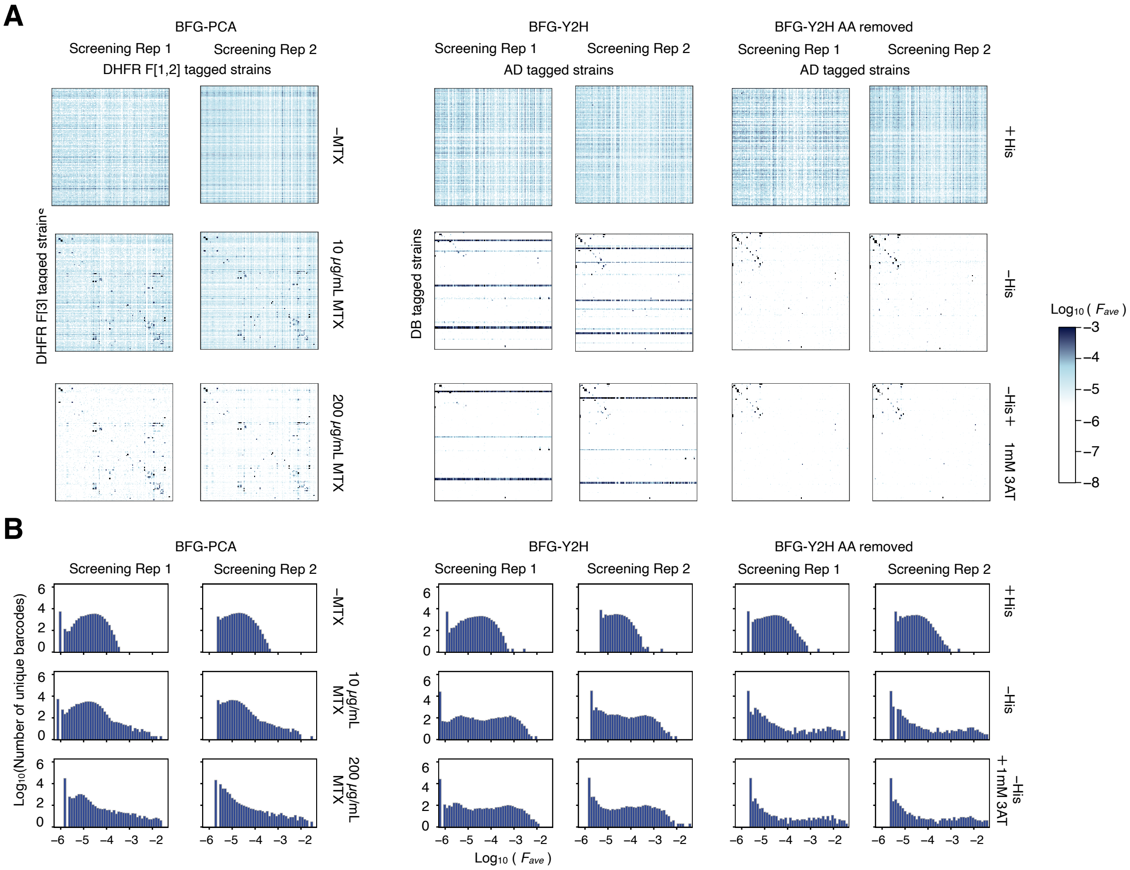


**Figure S5. Raw frequency of fused bait-prey barcodes in BFG-PCA and BFG-Y2H screenings.**

A. Heatmap of raw frequency (*F*) shown for each fused barcode in BFG-PCA and BFG-Y2H screenings. Note that the barcode fusion replicates (BC1-BC1 and BC2-BC2) were averaged for each bait-prey barcode. AA: Auto-activator.

B. Histogram representation of the bait-prey barcode abundance in each selection condition. AA: Auto-activator.

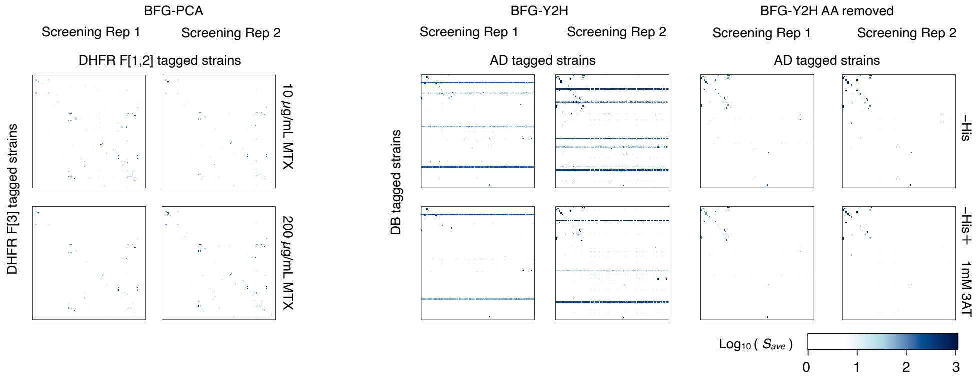


**Figure S6. Enrichment of bait-prey barcodes in selective conditions compared to their respective non-selection conditions.**

Heatmap of enrichment signal (*s*) shown for each bait-prey barcode in BFG-PCA and BFG-Y2H screenings. Note that the barcode fusion replicates (BC1-BC1 and BC2-BC2) were averaged for each bait-prey barcode. AA: Auto-activator.

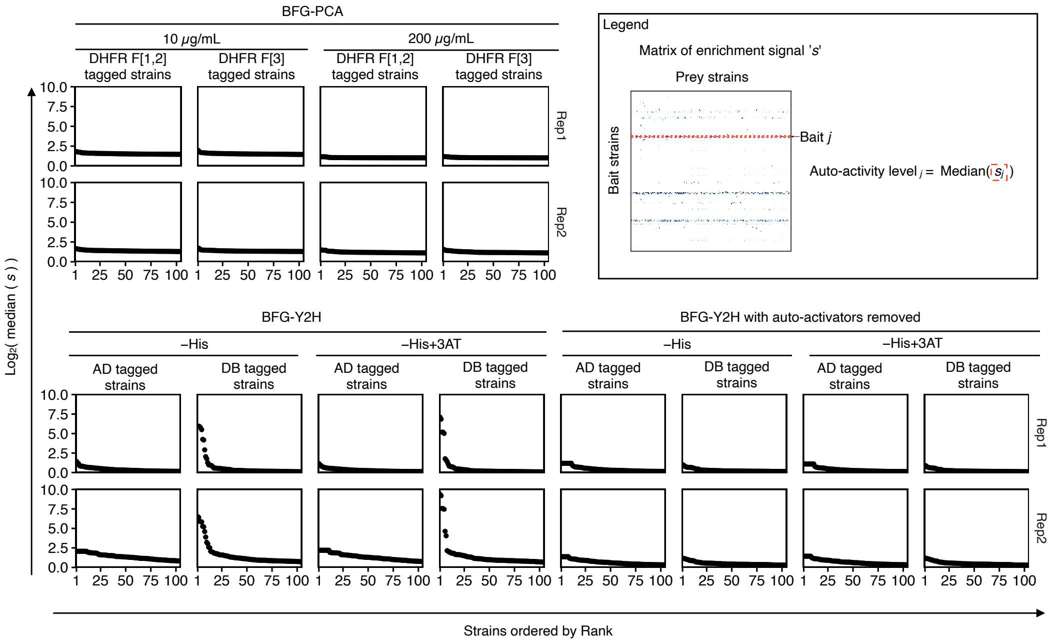


**Figure S7. Comparison of auto-activity on BFG-Y2H and BFG-PCA.**

Rank plot of bait/prey barcodes ordered by the median of enrichment score ‘s’. Some DB-X fused ORFs in Y2H screenings are known to have problematic auto-activation, where reporter activation occurs regardless of the interacting partner.

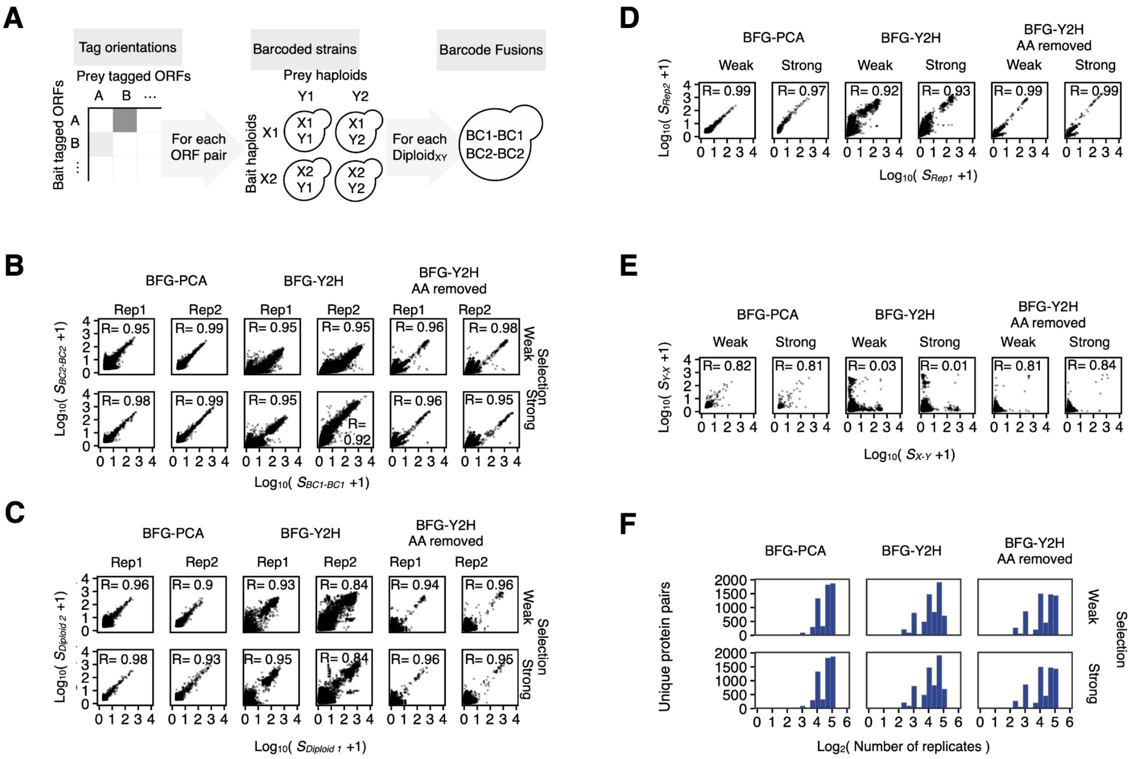


**Figure S8. Correlation of enrichment score between replicates.**

A. Schematic of the internal replicates within each screening. (Left) Protein pairs are tagged in two orientations, '*Bait-Prey*' and '*Prey-Bait*'. (Center) Each protein is assigned two barcode replicates. Four combinations of diploid strains, each having a unique bait-prey barcode, are expected upon mating. (Right) Each bait-prey barcode has two barcode fusion cassettes, BC1-BC1 and BC2-BC2.
B. Correlation of enrichment scores of BC1-BC1 and BC2-BC2 in each bait-prey barcode. Weak: -His or methotrexate 10μg/mL conditions according to methods. Strong: -His + 3AT or methotrexate 200 μg/mL conditions according to methods.
C. Correlation of enrichment scores of bait-prey barcode for each protein pair (orientation dependent). Diploids sharing any bait/prey barcodes were excluded from analysis.
D. Correlation of enrichment scores of screening replicates for each protein pair (orientation dependent). Barcode fusion and bait-prey barcode replicates were averaged.
E. Correlation of enrichment scores of screening orientation for each protein pair. Barcode fusion, bait-prey barcode, and screening replicates were averaged.
F. Histogram representing the total number of replicates per protein pair in BFG-PCA and BFG-Y2H screenings.

R: Pearson correlation coefficient. AA: Auto-activator.

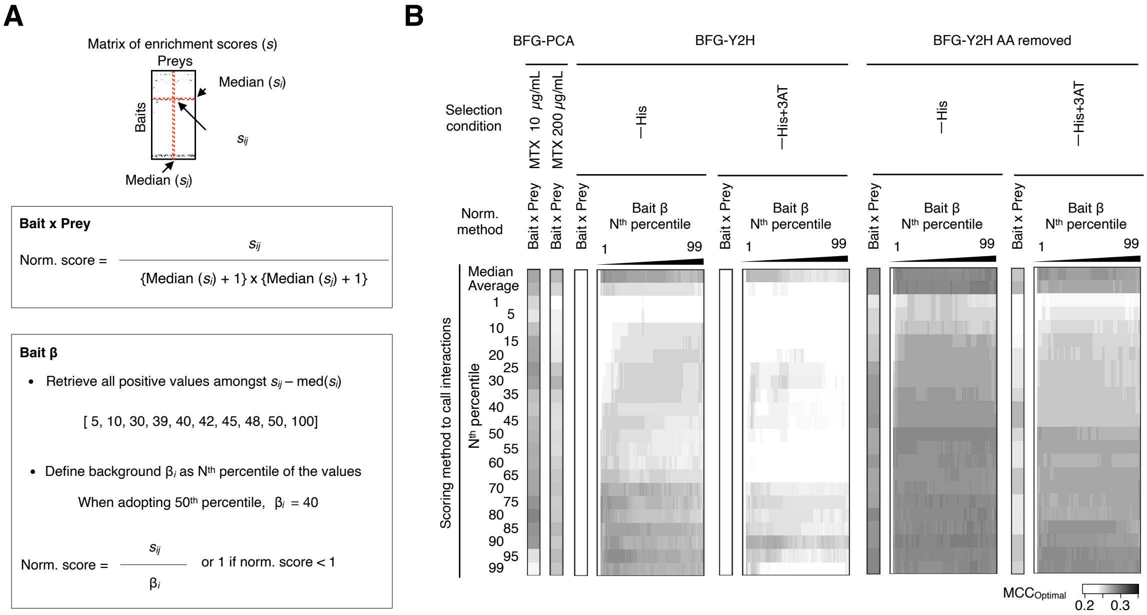


**Figure S9. Optimal MCC on normalization and scoring methods.**

A. Schematic of normalization of enrichment scores. (Bait x Prey) Since we observed both bait and prey barcodes exhibit backgrounds, we divided enrichment scores by the background of both bait and prey haploid strains. We added a constant of 1 to the background to reduce noise in the process. (Bait β) The method to normalize the BFG-Y2H score described in Yachie et al. Here the scores are normalized by dividing the enrichment score with the bait background level β, or 1 if the normalized score is below 1. Bait background level β is defined as the $N^{th}$ percentile of positive values of enrichment scores when subtracted by the median of the same bait strain.

B. Optimal MCC scores for each scoring method and normalization method tested. AA: Auto-activator.

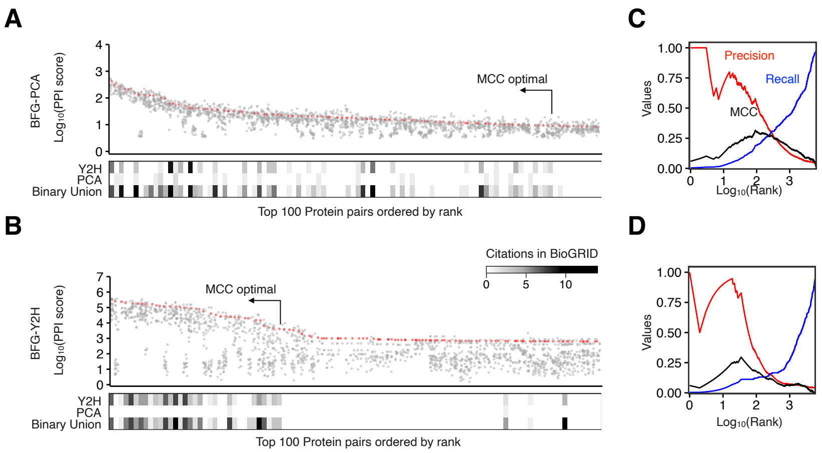


**Figure S10. Quality of BFG-PCA and BFG-Y2H screenings.**

A. & B. Result of assay ordered by PPI score rank for BFG-PCA (A), and BFG-Y2H (B). Grey dots represent replicates, and the red dots represent replicates on the percentile threshold used to call the ranks. The heatmap shows previously reported interactions in the BioGRID database. Binary union consists of interactions reported by Y2H, PCA, Biochemical activity, Affinity Capture-Luminescence, Reconstituted Complex, Co-crystal Structure, and FRET.

C. & D. Precision, recall, and MCC curve for BFG-PCA (C) and BFG-Y2H (D) with the given rank threshold.

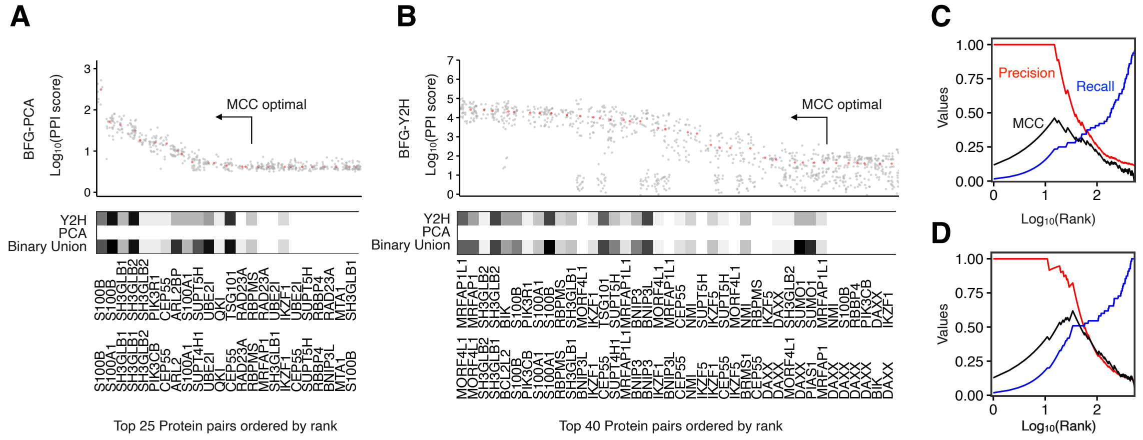


**Figure S11. Detected interactions in the Human protein subset.**

A. & B. Result of assay ordered by PPI score rank on the Human proteins for BFG-PCA (A)/BFG-Y2H (B). Grey dots represent replicates, and the red dots represent replicates on the percentile threshold used to call the ranks. The heatmap shows previously reported interactions in the BioGRID database. Binary union consists of interactions reported by Y2H, PCA, Biochemical activity, Affinity Capture-Luminescence, Reconstituted Complex, Co-crystal Structure, and FRET.

C. & D. Precision, recall, and MCC curve for BFG-PCA (C) and BFG-Y2H (D) on Human protein pairs.

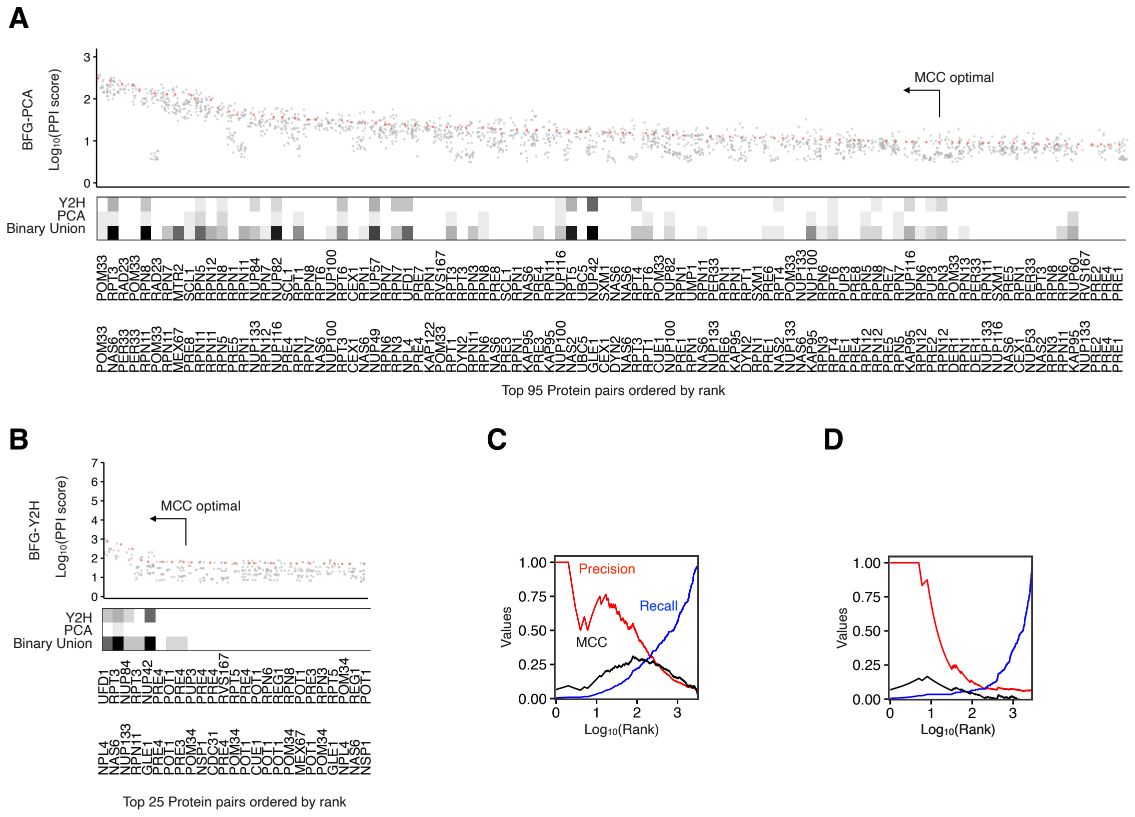


**Figure S12. Detected interactions in the Yeast protein subset.**

A. & B. Result of assay ordered by PPI score rank on the Yeast proteins for BFG-PCA (A)/BFG-Y2H (B). Grey dots represent replicates, and the red dots represent replicates on the percentile threshold used to call the ranks. The heatmap shows previously reported interactions in the BioGRID database. Binary union consists of interactions reported by Y2H, PCA, Biochemical activity, Affinity Capture-Luminescence, Reconstituted Complex, Co-crystal Structure, and FRET.

C. & D. Precision, recall, and MCC curve for BFG-PCA (C) and BFG-Y2H (D) on Yeast protein pairs.

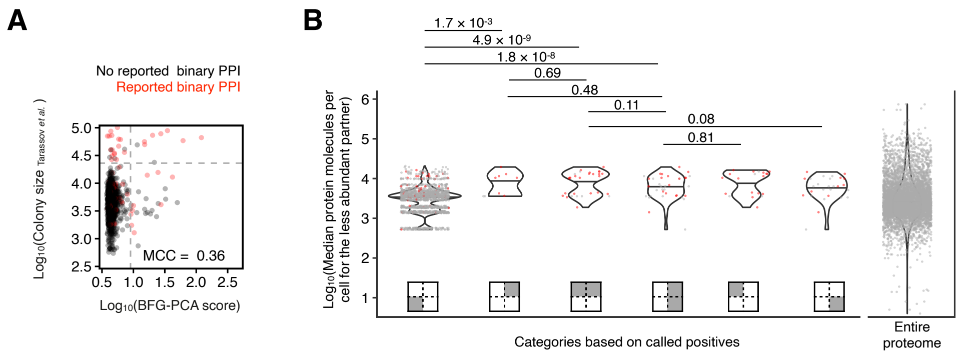


**Figure S13. BFG-PCA scores compared to previous DHFR-PCA datasets.**

A. Scatter plot of normalized BFG-PCA PPI score and the colony intensity data on methotrexate-selected plates from Tarassov et al. Grey dashed lines represent the threshold to call positives. Red represents PPIs in the BioGRID database reported by binary PPI detection methods.

B. Violin plot of protein molecules of the less abundant protein amongst the pair, categorized by sections of the scatterplot. Horizontal line represents the $50^{th}$ quantile of the data. For the PPI score data, red dots represent previously reported binary interactions. Statistical test: Mann Whitney U-test.

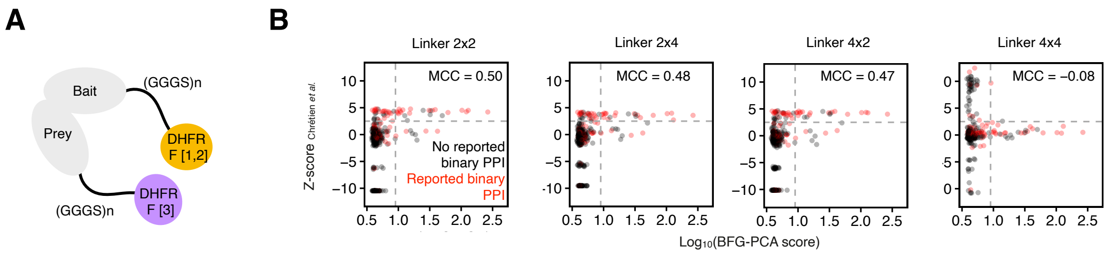


**Figure S14. BFG-PCA scores compared to DHFR-PCA with extended linkers.**

A. Illustration of DHFR-PCA linker design. Chrétien et al. detected PPIs with linkers with $(GGGS)_2$ or $(GGGS)_4$ in both bait and prey orientations.
B. Scatter plot of BFG-PCA score and Z-score obtained in Chrétien et al. Grey dashed lines represent the threshold to call positives. Red represents PPIs in the BioGRID database reported by binary PPI detection methods.
